# Global climate gradients structure soil *Legionella* diversity, relative abundance and pathogen distributions

**DOI:** 10.64898/2026.03.10.710744

**Authors:** Hans W. Singh, Mengting Maggie Yuan

## Abstract

Legionnaires’ disease has increased sixfold since 2000 in the United States, yet environmental reservoirs such as soils remain understudied sources of infection. It remains unclear how the diversity and abundance of environmental pathogens, such as *Legionella*, will shift as the climate continues to change. Here, we utilize 4,287 publicly available global soil 16S rRNA gene amplicon datasets to analyze how the distribution of the genus *Legionella*, including pathogenic lineages, varies across continental-scale biogeographic regions and along climatic and geochemical gradients. *Legionella* relative abundance increased across the joint precipitation-air temperature gradient, and mixed-effects modeling indicated that the chance of detecting *Legionella* in soils increased substantially when annual precipitation reached ∼500 mm. *Legionella* diversity was characterized by high spatial turnover, with the majority of abundant ASVs restricted to individual regions despite broad phylogenetic representation across the genus. Further, fewer than 2% of our 16S-derived *Legionella* sequences matched ASVs corresponding to characterized species, and the clinically dominant species *L. pneumophila* was rarely detected. In contrast, ASVs matching several other known pathogens (*L. longbeachae, L. anisa, L. bozemanae, L. cincinnatiensis*) were orders of magnitude more common than *L. pneumophila* in soils. Moreover, pathogenic *Legionella* ASVs were detected significantly more frequently at warmer and wetter conditions. Collectively, our findings suggest that *Legionella* relative abundance and diversity in soils, especially pathogenic lineages, are shaped by both dispersal and climatic filtering. As temperature and precipitation regimes shift in the future, our results imply that soils should be an important pathogen source to monitor and that *Legionella* species beyond *L. pneumophila* warrant increased ecological and public health attention.

## Introduction

The gammaproteobacterial genus *Legionella* is the causative agent of Legionnaires’ disease, a severe, pneumonia-like pulmonary infection that has increased sixfold since 2000 in the United States (1). Currently, the rate of Legionnaires’ disease is over 2.4 cases per 100,000 people in both the U.S. and Europe (2–3), and it carries a high economic burden ($800 million annually) (4), hospitalization rate (95%) and case fatality rate (10%) (2). *Legionella* can also cause the milder, flu-like Pontiac fever, and together these two illnesses are collectively referred to as legionellosis (1). *Legionella* has tight evolutionary relationships with free-living amoebae and other protozoan hosts, which function as analogous systems to human alveolar macrophages. Mechanistically, *Legionella* utilize type I, II, and IV secretion systems to deliver a large suite of effector proteins (∼300) that hijack host cellular functions and enable intracellular survival (1,5–8). While *Legionella* spp. have evolved the capacity to replicate intracellularly, they can also exist as free-living or within biofilms, and are commonly detected in freshwater systems (1,9). Despite the widespread environmental occurrence of *Legionella* (10), the climatic, geochemical, and ecological factors that shape their global distribution and diversity across natural reservoirs, such as soils, remain poorly understood.

Following its first identification associated with cooling towers (11), *Legionella* has been intensively studied as a human pathogen in built water systems (12), but the ecology of the genus in soil and freshwater remains far less clear. Typical *Legionella* infections follow inhalation of aerosolized water from contaminated built systems like cooling towers and showers (12), and usually involve *L. pneumophila* (80% of cases in the U.S. and Europe) (1–3). Direct exposure to *Legionella* from environmental reservoirs such as soils remains understudied as a source of legionellosis, although soil-associated *Legionella* have been shown to reach humans through aerosolization or hydrological transport (13). It is important to note that 95% of legionellosis cases in the U.S. are diagnosed using urinary antigen tests, which, while rapid and convenient, are limited to detecting *L. pneumophila* serogroup 1 (14–15). Notably, 23 of the 73 characterized *Legionella* species have been implicated in human infections, highlighting a significant gap in diagnostic sensitivity and species-level resolution (16–38). Legionellosis incidence over the past few decades is strongly correlated with precipitation, temperature and humidity (39–40). These observations are supported by increased infection rates in summer and fall months (2–3) when precipitation and temperature peak in most climate types. Sporadic, community-acquired cases without clear sources are also rising, with 66% of recent cases not linked to healthcare, travel or senior living (2). As cases grow despite improved disinfection and surveillance, environmental reservoirs beyond built systems should be considered as transmission sources. In Australasia, over half of reported cases are caused by *L. longbeachae*, a species traced to potting soils (1,41), implying a potentially significant contribution of infections not associated with built water systems.

*L. pneumophila* dominates genome sequencing efforts (79.7% of complete *Legionella* genomes in the NCBI database) (42), yet their well-studied physiology and pathogenesis do not necessarily apply to other species. For example, *L. longbeachae* uses a different intracellular mechanism and only shares 30% of its effector proteins with *L. pneumophila* (43–44). Effector diversity is high and largely species-specific, with eukaryotic-like domains commonly seen across the genus, reflecting cases of gene acquisition from their hosts (45). While *Legionella* effectors constitute a taxonomically diverse virulence factor set, implying varied host interactions, most remain poorly understood (46). As non-*pneumophila* infections are likely underrecognized, their risks under global change are highly uncertain. It remains unclear how the diversity and abundance of environmental pathogens, including *Legionella*, will shift as the global climate continues to change. Environmental reservoirs such as soils may be significant yet underrecognized sources of infection. Soils in particular remain poorly represented in culture-based and genomic surveys of *Legionella* diversity, limiting our understanding of legionellosis risk (46–47). Understanding the global diversity and relative abundance of *Legionella* species is an essential first step to assess their potential links to the rising incidence of legionellosis.

Over the past two decades, the acceleration in genomic, metagenomic, and high-throughput amplicon sequencing capabilities have revolutionized our ability to perform large-scale assessments of microbial diversity (48–50). In conjunction, the development of large data repositories such as NCBI and JGI has enabled the accumulation of microbiological datasets that are often linked to accompanying metadata (42,51). From a sequencing perspective, *Legionella* is a rare taxon in natural environmental systems, making it difficult to recover in metagenomic bins. For example, of the nearly 350,000 high-quality and medium-quality environmental metagenomic bins within the JGI database, only 109 (0.03%) belong to the genus *Legionella*, highlighting the challenges of using metagenomic sequencing to assess the global diversity of rare taxa (51). Complete genomes across a breadth of strains can also be challenging to compare due to biases in target samples (towards infected patients) and cultivation techniques (towards *L. pneumophila*-specific media) (15,42,47). High-throughput targeted amplicon sequencing of published *Legionella*-specific primers is also insufficient for a global comparison of *Legionella* diversity as they are rarely utilized over large geographic distances, primers are biased towards clinical species, and infrequently targeted in soil samples (52). The most widely used 16S rRNA gene primer set to date (515F-806R) optimizes broad prokaryotic taxonomic coverage and is utilized in large-scale analysis of microbial diversity across every continent (12,48). These attributes make it optimal for a global analysis, with large numbers of samples and high sequencing depth compensating for the rarity of *Legionella* reads.

Nonetheless, taxonomic classification of 16S reads to rare members of the microbiome remains a challenge (53–54). Utilization of both k-mer based and identity-based classifiers depend on existing reference sequences, but the sparse and *L. pneumophila*-skewed representation of *Legionella* in 16S databases such as SILVA, Greengenes and GDTB limits accurate detection (42,55–56). When a query sequence lacks a close match in the database, classifiers default to higher taxonomic levels (e.g., class or order), thereby masking genus- or species-level diversity (54–55). A further limitation within the sequencing pipeline is that rare taxa like *Legionella* often resolve into singleton amplicon sequence variants (ASVs), which are often removed during quality filtering (57–58). Phylogenetic insertion offers the most accurate approach for assigning taxonomy from short 16S amplicons, particularly for rare taxa, as they place sequences on an evolutionary backbone rather than relying on k-mer frequencies or pairwise identity (59–61). These methods are computationally intensive and impractical for analysis of complete 16S datasets (59), but become more tractable when applied to targeted analyses of a single genus, such as *Legionella*.

Here, we analyze the global soil ecology of *Legionella* using 4,287 publicly available 16S rRNA amplicon datasets spanning diverse biogeographic regions and environmental gradients. Specifically, we identify environmental predictors of *Legionella* abundance, characterize patterns of diversity and biogeographic turnover, and assess the prevalence of ASVs corresponding to known pathogenic *Legionella* species.

## Results

### Two-step taxonomic approach accurately classifies *Legionella* sequences

To maximize the recovery, speed and accuracy of *Legionella* detection, we implemented a two-step taxonomic classification pipeline in which 16S amplicon sequences (the first 150 bp following the 515F primer) were first screened against a custom BLAST search, after which candidate *Legionella* hits were inserted into a backbone reference phylogeny built from representative and complete genomes. Analysis using all Gammaproteobacteria sequences within the Silva database highlights the accuracy of our approach - every *Legionella* sequence (n=1586) in SILVA was correctly detected and placed within the *Legionella* clade, and no non-*Legionella* Gammaproteobacteria sequences (n=16,926) were phylogenetically verified as *Legionella* sequences (i.e., no false positives). As an additional validation to our pipeline, we sequenced a mock community consisting of DNA from *L. pneumophila*, *L. longbeachae* and *L. micdadei*, and as expected found these to be the only three ASVs recovered.

### Relative abundance of *Legionella* sequences across climatic and biogeochemical gradients

In total, 4287 globally distributed soil 16S rRNA amplicon sequencing samples were analyzed, representing eleven distinct BioProjects (58–73) across eight distinct geographic locations: North America, Europe, Africa, Australia, Hawaii, Iceland, China, and Chile (Fig. 1a-b). The bioprojects we selected broadly fell into one of two categories, with the first comprising microbial surveys carried out across large biogeographical ranges that span substantial climatic and biogeochemical gradients. Our second category corresponded to bioprojects from fixed geographic locations that imposed controlled environmental manipulations: temperature in Iceland, precipitation in Iowa and both temperature and precipitation in Oklahoma. Across our entire dataset, the average sequencing depth was 56,960 sequences with 22.76 *Legionella* sequences recovered (0.039%). Despite *Legionella* being a rare taxon, the overwhelming majority of our samples (93.0%) contained at least one *Legionella* read, highlighting that *Legionella* are a consistent member of soil microbial communities.

**Fig. 1.**
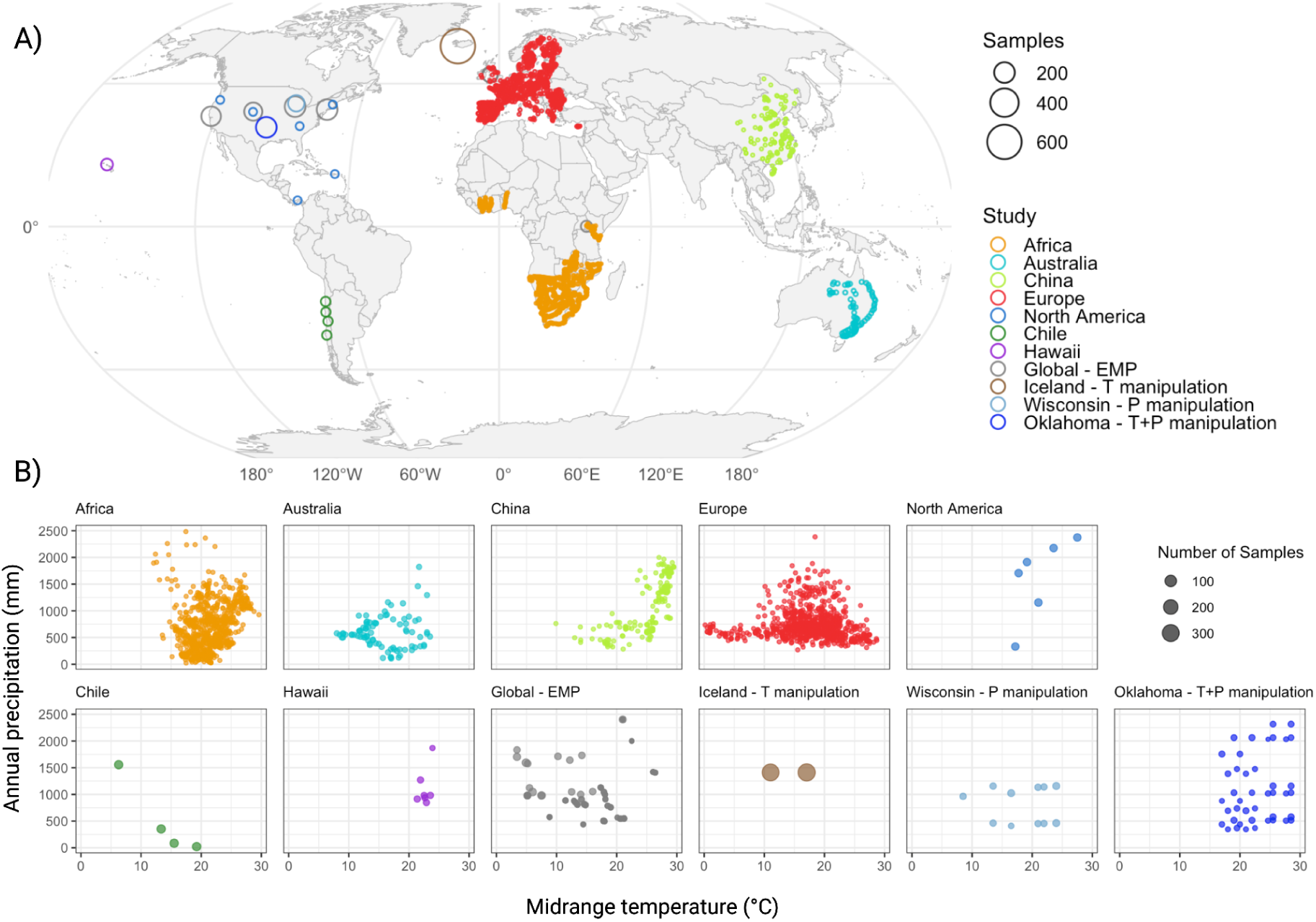
Samples selected for global analysis. A) Geographic distribution of soil samples used in this study. Colors indicate sequencing projects and point size corresponds to the number of samples at each location. B) Climatic ranges for each sequencing project. Points show individual sampling locations plotted by midrange temperature (°C) during the sampling month (x-axis) and annual precipitation (mm) (y-axis). Panels correspond to sequencing projects, and point size indicates number of samples.

From each sample, the following metadata were collected as indicators of the climatic and biogeochemical conditions that *Legionella* was recovered from: midrange monthly temperature, annual precipitation, soil pH, total organic carbon, and total organic nitrogen. When we analyzed how *Legionella* relative abundance varied across climatic and biogeochemical variables, Spearman correlations revealed weak but significant monotonic relationships (Fig. S1) with all environmental variables (p<10^-8^). The strongest associations were negative with elevation (ρ = –0.24) and positive with soil carbon content (ρ = 0.24). Annual precipitation (ρ = 0.2) and soil nitrogen content (ρ = 0.18) displayed moderate positive correlations, while pH (ρ = –0.13) and temperature (ρ = 0.09) exhibited weaker correlations. Although statistically robust due to large sample sizes, all correlations were small in magnitude, indicating that *Legionella* abundance is only weakly structured by any single climatic or biogeochemical gradient.

Since Spearman correlations only quantify monotonic trends and do not capture non-linear or threshold responses, we next evaluated differences in *Legionella* relative abundance across discrete environmental categories (Fig. 2a, Fig. S2). Each environmental variable was partitioned into five buckets representing the full range of metadata, and Kruskal-Wallis tests revealed highly significant differences among buckets for all six variables (all p < 10^-15^). Dunn’s post hoc comparisons show clear, structured shifts in abundance across these buckets. For temperature, *Legionella* abundance was significantly higher in the 25-30°C range compared to all other temperature ranges (Fig. 2a). Precipitation showed a strong threshold response: the driest sites (0-400 mm) had significantly lower abundance than the other four wetter categories, which formed a single statistically homogeneous group (Fig. 2a). Soil carbon exhibited a graded increase in abundance with each successive bucket (0-1% < 1-2%, 2-3% < 3-4% < 4+%), producing four distinct groups (a-d) (Fig. S2). Nitrogen content followed a similar positive pattern with low-N soils (0-0.1%), moderate-N (0.1-0.2%, 0.2-0.3%, 0.3-0.4%) and high-N (0.4+%) soils falling into three distinct groups (a-c) (Fig. S2). For pH, abundance was highest under strongly acidic conditions (<5), decreased across intermediate pH levels (5–8) and was lowest in alkaline soils (8+), forming three groups (a-c) (Fig. S2). Lastly, elevation strongly structured *Legionella* abundance, which displayed a clear low elevation optimum (0-200 m.), declined at middle elevations (200-400 m.) and displayed minimum abundance at higher elevations (400-600 m., 600-800 m., 800+ m.) (Fig. S2). Collectively, these categorical patterns reveal a mix of gradual monotonic and threshold responses, suggesting that depending on the environmental driver, *Legionella* abundance can be shaped by both sharp environmental transitions and smooth continuous gradients.

**Fig. 2.**
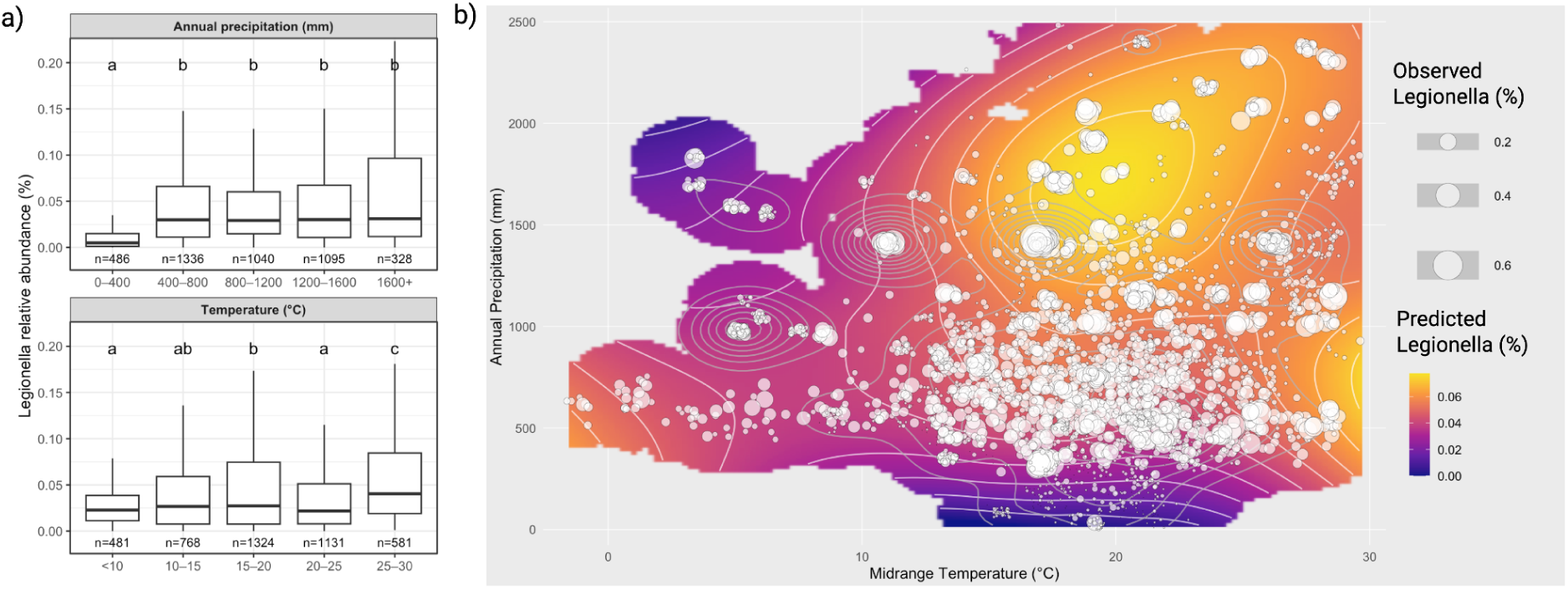
Climatic controls on *Legionella* relative abundance. (a) Relative abundance of *Legionella* sequences across climatic ranges. Samples are grouped into five bins for annual precipitation (upper panel) and midrange temperature during the sampling month (lower panel). Boxplots show the distribution of *Legionella* relative abundance within each bin. Letters indicate statistically significant differences among groups (Kruskal-Wallis with Dunn’s post hoc test). (b) Relative abundance of *Legionella* sequences across the joint temperature-precipitation gradient. Circles indicate observed *Legionella* relative abundance. The colored surface represents LOESS-predicted *Legionella* relative abundance, with contour lines indicating predicted abundance levels.

As climatic data are the easiest to monitor at scale, we next queried the interactive effects of climate on *Legionella* relative abundance by analyzing the two-dimensional environmental space of temperature and precipitation. A bivariate temperature-precipitation response surface revealed clear interactive structure, with the highest predicted *Legionella* relative abundances occurring under both warm conditions and high precipitation (Fig. 2b). A two-dimensional GAM (generalized additive model) revealed a highly significant joint temperature-precipitation effect (EDF = 21.7, p<10^-15^) and significantly improved model fit over a model with independent smooth functions for temperature and precipitation (ANOVA: F=2.82, p=0.012), confirming that *Legionella* relative abundance is shaped by nonlinear interactive effects of climatic conditions.

To evaluate the joint effects of all climatic and biogeochemical variables while accounting for site-to-site variation, we used a hurdle mixed-effects modeling framework, consisting of a binomial mixed model for presence-absence and a beta mixed-effects regression for relative abundance conditional on presence, with study included as a random intercept. Detection probability of *Legionella* sequences increased significantly with only annual precipitation (Fig. S3), whereas no other variables were associated with changes in presence of *Legionella*. In contrast, conditional relative abundance increased significantly with temperature and soil carbon content, and declined with increasing pH and soil nitrogen content (Fig. S3). Notably, annual precipitation showed no association with relative abundance. Together, these results indicate that moisture availability primarily governs *Legionella* occurrence, while temperature and local biogeochemical variables govern *Legionella* relative abundance. Further, holding all other climatic and biogeochemical variables constant and accounting for study-to-study variation altered the inferred effects of certain predictors, such as nitrogen content (positive in isolation, negative in joint model) and precipitation (positive in isolation, neutral in joint model).

### *Legionella* relative abundance at experimentally manipulated sites

Experimental field manipulations are powerful tools for identifying predictors of *Legionella* abundance, as they selectively alter target variables while keeping all other variables constant. Analysis of 16S data from a geothermally-warmed site in Iceland (+6°C) showed that *Legionella* relative abundance was significantly higher than in neighboring ambient sites (p<0.01) (Fig. S4). Similarly, analysis of rainfall-manipulation sites in Wisconsin revealed that *Legionella* increased with volumetric water content (r = 0.39, p < 0.01) (Fig. S5). Finally, at a site combining precipitation and temperature manipulations in Oklahoma, *Legionella* relative abundance was positively correlated with annual moisture (r = 0.32, p < 0.001) and negatively correlated with annual temperature (r = -0.34, p < 0.001) (Fig. S6). These results highlight the role of environmental selection in *Legionella* distribution.

### *Legionella* diversity across biogeographic, climatic and biogeochemical gradients

Non-metric multidimensional scaling (NMDS) based on Bray-Curtis dissimilarities revealed weak but significant regional structuring of *Legionella* community composition (Fig. S7). Although global samples largely overlapped in ordination space, Chilean samples exhibited a pronounced shift along NMDS1 and greater dispersion than other regions, indicating elevated within-region heterogeneity. Permutation-based multivariate analysis of variance (PERMANOVA) confirmed a significant effect of region on *Legionella* community composition (r^2^ = 0.02, p < 0.001). Permutation-based fitting of environmental variables onto the NMDS ordination identified pH as the only factor significantly correlated with Legionella community composition at the global scale (envfit, r^2^ = 0.0043, p = 0.043) (Fig. S8). While other environmental factors were not significant predictors, binned ordinations suggest gradual, directional turnover in community composition across the midrange temperature, soil carbon and soil nitrogen gradients (Fig. S8). For these variables, lower-value bins transition directionally across ordination space toward higher-value bins rather than forming discrete clusters, which is consistent with continuous, non-threshold responses to environmental variation.

To further resolve regional structuring within the dominant members of the *Legionella* community, we examined the 300 most abundant *Legionella* ASVs within each of our 8 regions. The majority of dominant *Legionella* sequences (70.0%) were only detected in one region, with just 11.5% observed in 4+ regions and none detected in all 8 regions (Fig. 3a). The proportion of region-restricted dominant taxa varied among regions, with isolated regions such as Chile (84.6%), Hawaii (81.3%) and Iceland (74.0%) displaying high levels of regional specificity, whereas Europe (49.3%) and North America (59.0%) showed less (Fig. 3a). Pairwise comparisons showed limited overlap between Chile and any other region (1-8%), and limited overlap was also observed for paired regions with vastly different climates: Iceland/Africa (4%), Iceland/Hawaii (4%) and Iceland/Australia (6%) (Fig. S9). Regions with less geographic distance or similar climates contained the highest overlap - North America/Europe (30%), Iceland/Europe (20%) and China/Europe (18%) (Fig. S9). Indeed, mapping climatic differences onto these pairwise patterns revealed that overlap in dominant *Legionella* ASVs declined with increasing environmental dissimilarity (Fig. 3b), with the percentage of shared ASVs negatively correlated with increasing differences in both annual precipitation (Spearman ρ = -0.45, p=0.017) and temperature (ρ = -0.37, p=0.05) between regions. Collectively, these results suggest that both geographic separation and climatic similarity are associated with the structure of dominant *Legionella* ASVs across regions.

**Fig. 3.**
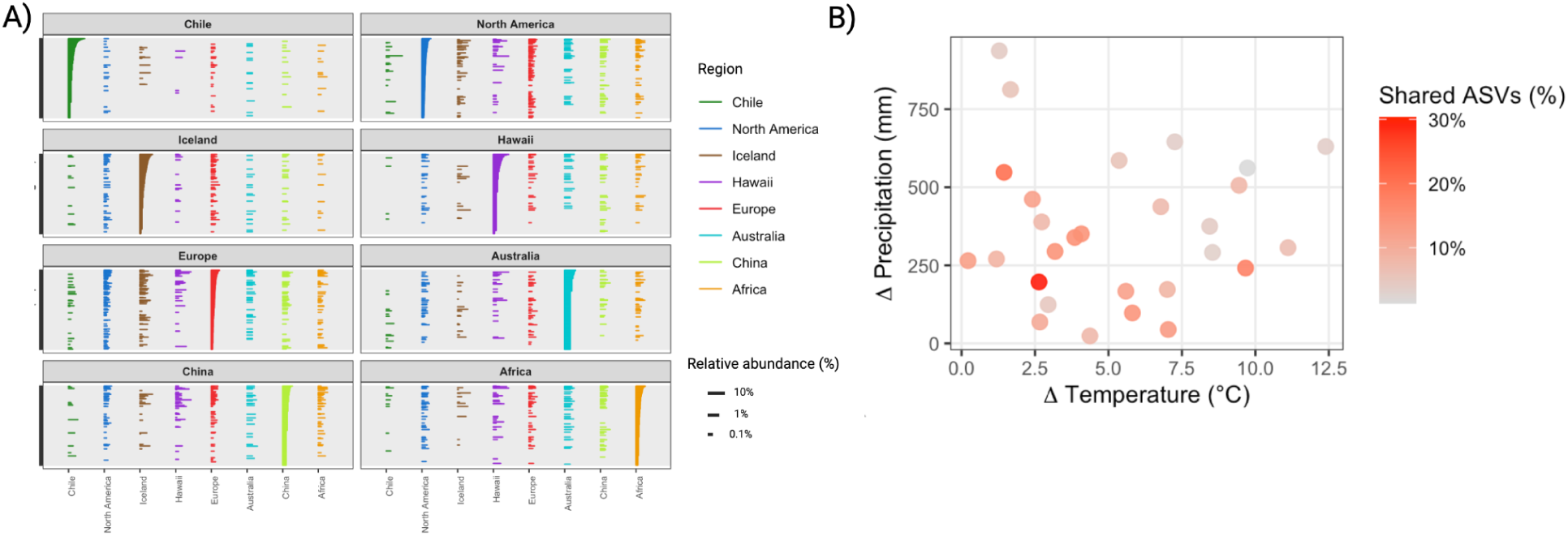
Regional distribution and climatic overlap of dominant *Legionella* ASVs. (a) Distribution of the 300 most abundant *Legionella* ASVs for each region. Each panel shows the dominant ASVs from one region, with the length of the bar indicating relative abundance, and colors indicating the presence of those ASVs across all 8 regions. (b) Climatic similarity and overlap of dominant *Legionella* ASVs among regions. Each point represents a pairwise comparison between regions, with axes showing differences in midrange temperature (°C) and annual precipitation (mm). Point color indicates the percentage of shared dominant ASVs between regions.

Finally, to determine whether the climatic and biogeographic-associated turnover in dominant *Legionella* ASVs corresponded to deep phylogenetic divergence, we constructed a maximum-likelihood phylogeny using IQ-TREE based on the 300 most abundant ASVs from each of the eight regions. Environmental ASVs spanned multiple deeply branching lineages and were broadly interspersed with both cultured and pathogenic *Legionella* reference sequences, indicating that abundant environmental *Legionella* represent a wide range of evolutionary lineages rather than a restricted subset (Fig. 4). Pathogenic *Legionella* references were distributed across multiple phylogenetic lineages rather than forming a single monophyletic group, consistent with phylogenetically widespread pathogenic potential within the genus. Although ASVs from different biogeographic regions did not segregate into discrete, region-specific clades and instead co-occurred throughout the phylogeny, null-model analysis revealed significant phylogenetic structure within several regions. Net Relatedness Index (NRI) values indicated significant deep phylogenetic clustering in North America, Iceland, Europe, China, and Africa, whereas Chile alone (lowest average midrange temperature and annual precipitation) exhibited strong deep overdispersion (Fig. S10). Nearest Taxon Index (NTI) values further revealed significant tip-level clustering in most regions, including Europe, Iceland, Africa, North America, Australia, and Chile (Fig. S10). These results indicate that while dominant environmental *Legionella* lineages are broadly distributed across the genus, climatic and biogeographic filtering operate within regions to structure communities at both deep (NRI) and fine (NTI) phylogenetic scales, with substantial heterogeneity among regions.

**Fig. 4.**
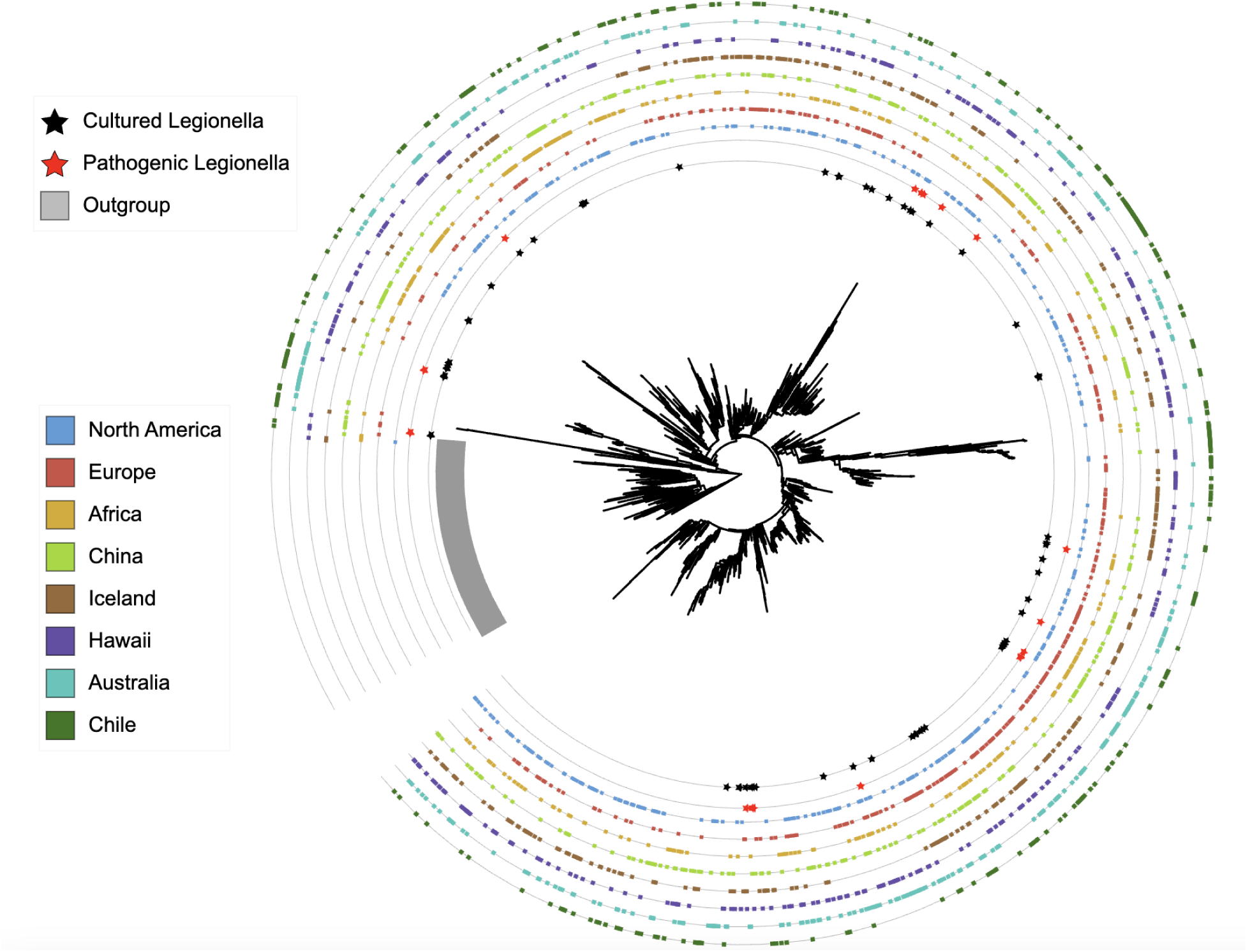
Phylogeny of dominant *Legionella* ASVs. The 300 most abundant *Legionella* ASVs from each region were placed into a maximum-likelihood phylogeny. Stars on the innermost rings indicate ASVs matching representative or complete genomes of cultured *Legionella*, with red stars marking human pathogens. Colored outer rings indicate the presence of each ASV among the dominant *Legionella* ASVs in each region.

### Spatial structuring of *Legionella* diversity

To evaluate spatial structuring of *Legionella* community composition at a finer scale, we quantified pairwise beta diversity among samples using Bray-Curtis dissimilarity calculated from *Legionella*-only ASV profiles and examined its relationship with geographic distance. Community similarity declined systematically with increasing geographic distance, exhibiting a significant distance-decay relationship (Fig. S11). Fitting a power-law model to our data demonstrated a strong negative scaling between similarity and distance (β = -0.134, p < 0.001), indicating that *Legionella* communities become progressively more dissimilar as spatial separation increases. Multiple regression on distance matrices (MRM) indicated that geographic distance remained a significant predictor of community dissimilarity after accounting for climatic and biogeochemical differences among sites (p < 0.001).

### Community context of *Legionella*-rich microbiomes

To identify microbial community features associated with *Legionella*-rich samples, we performed a differential abundance analysis using 97% OTUs. Samples were stratified into upper (>0.062%) and lower (<0.009%) quartiles of *Legionella* relative abundance, and in total 114 OTUs were enriched in *Legionella*-rich samples and 290 OTUs were enriched in *Legionella*-poor samples (|log₂ fold change| ≥ 2.5, FDR < 0.01) (Fig. S12). OTUs enriched in *Legionella*-rich samples were most frequently classified as Bacteroidota (35.1%) and Acidobacteria (27.2%) (Fig. S12). In contrast, the sharpest compositional shift was observed for Actinomycetota, which comprised 30.7% of OTUs enriched in *Legionella*-poor samples but only 3.5% of OTUs enriched in *Legionella*-rich samples (Fig. S12). Notably, many Actinomycetota OTUs enriched in *Legionella*-poor samples were classified to the family Rubrobacteria (33.7%), a lineage commonly associated with arid soil environments. Differentially abundant OTUs were broadly prevalent within their respective groups, with median prevalence of 40.7% among OTUs enriched in *Legionella*-poor samples and 38.4% among OTUs enriched in *Legionella*-rich samples, indicating community-wide shifts rather than effects driven by rare taxa.

### Relative abundance and diversity of *Legionella* ASVs from known pathogens

In total, 257 *Legionella* genomes in the NCBI database are annotated as either reference-quality or complete, representing 74 distinct species. When we clustered the first 150 bp following the 515F forward primer, hereafter referred to as the target 16S region, we found that this region resolves 51 species into unique (non-shared) ASVs (Fig. 5a). Only six ASVs were shared among multiple species, with one ASV representing twelve different species - *L. anisa*, *L. cherii*, *L. dumoffii*, *L. gormanii*, *L. longbeachae*, *L. parisiensis*, *L. qingyii*, *L. resiliens*, *L. santicrucis*, *L. sheltonii*, *L. sp 227* and *L. sp PC1000* (Fig. 5a). Of the 23 recognized pathogenic *Legionella* species, 12 were represented by unique ASVs (known pathogen ASVs), whereas the remaining 11 clustered into ASVs shared with at least one species not confirmed to be pathogenic (potentially pathogenic ASVs) (Fig. 5a). It is important to note that *Legionella* species are designated as pathogenic based on culture-confirmed human infections, a criterion that almost certainly underestimates pathogenic potential across the genus.

**Fig. 5.**
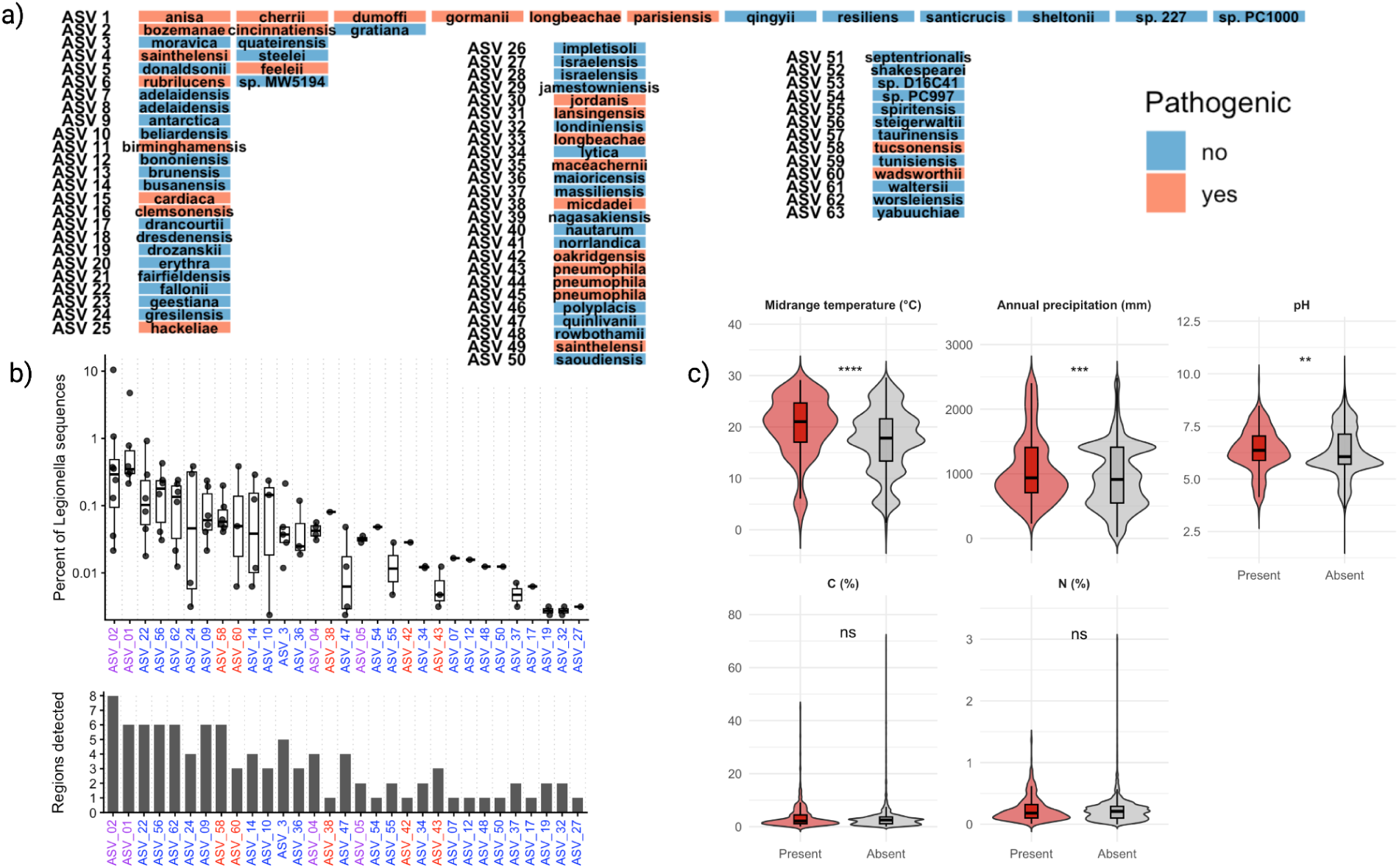
Distribution and environmental associations of characterized *Legionella* ASVs. (a) Clustering of characterized *Legionella* species based on the first 150 bp following the 515F primer of the 16S rRNA gene. Species sharing identical sequences are grouped into the same ASV and colored according to whether the species is a known human pathogen. (b) Relative abundance and geographic distribution of characterized *Legionella* ASVs. The upper panel shows the percentage of total *Legionella* sequences in each region that mapped to a given characterized ASV. Points represent individual regions and boxplots summarize the distribution across regions where the ASV was detected. X-axis labels are colored by pathogenic status (red = pathogenic, purple = potentially pathogenic, blue = non-pathogenic). ASV numbers correspond to those shown in panel (a). The lower section of panel (b) shows the number of regions in which each ASV was detected. (c) Environmental distribution of pathogenic or potentially pathogenic *Legionella* ASVs. Violin plots show the distribution of samples in which pathogenic ASVs were present (red) or absent (grey) across climatic and biogeochemical variables.

When we searched our dataset for pathogenic or potentially pathogenic *Legionella* ASVs (pASVs), the most abundant pASV matched two pathogenic species (*L. bozemanae* and *L. cincinnatiensis*), whereas the next most abundant pASV matched six pathogenic species (*L. anisa*, *L. cherrii*, *L. dumoffii*, *L. gormanii*, *L. longbeachae* and *L. parisiensis*), accounting for 0.48% and 0.43% of total *Legionella* sequences, respectively (Fig. 5b). Across all *Legionella* ASVs from our global analysis, these two pASVs ranked as the second and fourth most abundant sequences overall (Fig. S13). Consistent with their high global abundance, both pASVs were consistently detected across six of our eight geographic regions (>0.1% of *Legionella* sequences), with Chile and Iceland as exceptions. The only other pathogenic or potentially pathogenic *Legionella* ASVs recovered were those matching to *L. tucsonensis* (0.11%), *L. sainthelensi* (0.03%), *L. micdadei* (0.03%), *L. feeleii* (0.02%), *L. oakridgensis* (0.01%), *L. wadsworthii* (0.01%) and *L. pneumophila* (0.01%) (Fig. S5b).

To evaluate whether the presence of pathogenic *Legionella* was associated with environmental gradients while accounting for detection bias, we compared samples in which *Legionella* pASVs were present or absent across our environmental variables and fitted a logistic regression model including sequencing depth as a covariate. Our results illustrate that pASVs tended to occur significantly more often at higher midrange temperatures and under wetter climate conditions (Fig. 5c). Consistent with these patterns, logistic regression revealed that after controlling for sequencing depth, both midrange temperature and annual precipitation remained significant predictors of *Legionella* pASVs (p < 10^-5^). Model coefficients indicate that a 10°C increase in midrange temperature was associated with a 55% increase in the odds of detecting pathogenic *Legionella* ASVs, and a 500 mm increase in precipitation was associated with an approximately 45% increase in detection odds. In contrast, pH, soil carbon, and nitrogen content were not significant predictors of pathogenic *Legionella* ASVs after accounting for sequencing depth.

## Discussion

By leveraging 4,287 publicly available global soil 16S rRNA datasets coupled with an efficient rara-taxa detection method based on targeted phylogenetic insertion, we found that the rare (only 0.039% of the overall soil microbiome) but widespread (present in 93% of soil samples) gammaproteobacterial genus *Legionella* exhibits predictable environmental and biogeographic patterns across global soils. Warmer and wetter environments were consistently associated with higher relative abundances of *Legionella* and increased probabilities of detecting pathogenic *Legionella* ASVs, highlighting the role of climate in structuring environmental pathogen reservoirs. Consistent with these patterns, *Legionella* community composition also exhibited continental-scale structuring, with dominant ASVs showing strong geographic specificity and decreasing overlap among regions with more distinct climates. Notably, the most clinically significant species, *L. pneumophila* (1), was rare in soils, whereas the soil-associated *L. longbeachae* was expectedly among the most abundant *Legionella* ASVs detected, providing empirical support for the long-standing view that different pathogenic *Legionella* species occupy distinct environmental reservoirs (41). Together, these findings suggest that *Legionella* persistence in soils is governed by a set of ecological filters: geographic distance and climate influences which lineages dominate regionally within the global species pool; soil water availability determines establishment, and temperature and local biogeochemical conditions regulate population growth.

### Climatic influence on *Legionella* presence and relative abundance in global soils

Mapping the ecological niches of environmental pathogens is increasingly important as climate change alters environmental conditions. The combination of extremely low relative abundance yet near-ubiquitous detection across soils suggests that *Legionella* behave as a member of the microbial “rare biosphere” (74), persisting at low abundance across diverse environments while responding opportunistically to favorable ecological niches. Such regulation of microbial growth and activity often reflects environmental controls, particularly by temperature and precipitation, which are strong drivers of the distribution and transmission of many human pathogens–from dengue viruses to *Leptospira* bacteria (75–76). Temperature governs biochemical process rates and often exerts selective pressure on human pathogens; precipitation influences air humidity, soil hydrology, solute movement, and nutrient deposition, which are all critical controls of microbial activity in the air, soil, and aquatic systems (77). In addition, surface air temperature and precipitation are regularly monitored globally with a high spatial resolution (78), and many of these datasets are publicly available, providing a pragmatic framework for disease risk assessment at scale.

Previous work showed that Legionnaires’ disease incidence is positively associated with warmer and wetter climate (79); our finding that *Legionella* relative abundance increased globally in soils along the joint temperature-precipitation gradient implies that increasing environmental load under these conditions may contribute to exposure and infection. Warmer and wetter soils may favor *Legionella* persistence through several mechanisms. Elevated soil moisture promotes biofilm formation and supports higher densities of free-living amoebae and other protist hosts that facilitate intracellular replication (9,72), while warmer temperatures increase microbial activity and host-pathogen interactions (80–82). Consistent with this biofilm-and host-mediated ecology, our mixed-effects modeling identified precipitation as the primary determinant of *Legionella* presence. In contrast, temperature and local biogeochemical variables were more strongly associated with changes in *Legionella* relative abundance, suggesting that different environmental factors govern establishment and population growth. This pattern is consistent with general principles of soil microbial ecology, in which moisture availability constrains habitat suitability while temperature and nutrient availability regulate metabolic rates and microbial growth (83–84). Notably, a previous global analysis showed that relative abundance of the order *Legionellales*, including multiple genera within the families *Legionellaceae* and *Coxiellaceae*, decreases with soil temperature (10). The discrepancy between this previous study and ours likely reflects differences in taxonomic resolution (order versus genus) as well as precipitation effects that were not considered in the previous analysis. As sporadic, community-acquired cases of legionellosis continue to rise (2), our findings collectively reinforce the possibility that environmental reservoirs such as soils may represent significant yet underrecognized sources of *Legionella* exposure.

Patterns in the broader soil microbiome further support the ecological context of *Legionella* persistence. *Legionella*-rich samples were enriched in Bacteroidota and Acidobacteria, groups commonly associated with soils containing higher organic matter availability and lower pH, respectively (85–86). This pattern aligns with our observation that *Legionella* relative abundance was higher in carbon-rich and more acidic soils. In contrast, Actinomycetota were strongly enriched in *Legionella*-poor samples; these taxa are commonly associated with drier soils and are well known for producing antimicrobial secondary metabolites (87–88), which may inhibit the presence of other microbial taxa, including *Legionella*. Under culture conditions, Pseudomonas has been shown to suppress *Legionella* growth through antimicrobial metabolites and quorum-sensing molecules (89), supporting the idea that microbial antagonism may also affect *Legionella* persistence in natural environments. However, bacterial-bacterial interactions involving *Legionella* in soil systems remain poorly characterized. Understanding interactions between *Legionella* and surrounding microbial communities may therefore be critical for determining the ecological conditions that promote or suppress its persistence, including potential competitive suppression, facilitation within biofilms, or indirect effects mediated through protist hosts.

The rapid expansion of sequencing technology and deposition of microbiome datasets in public repositories have enabled large-scale global analyses that were previously infeasible to conduct (90–91). However, as with many data reuse studies, this approach introduces potential biases, including uneven geographic sampling, variable sequencing depth and study-to-study methodological variation. Nonetheless, the large number of samples analyzed, consistency of these patterns with experimental field manipulations, and the use of hurdle mixed-effects models accounting for sequencing depth and study-level variation suggest that the broad climatic patterns observed here are robust. Although our analysis is based on relative abundance, previous studies have shown that warmer and wetter soils generally support higher microbial activity (81,83), suggesting that increases in *Legionella* relative abundance observed here may also correspond to elevated absolute abundance. However, confirming this will require future work using *Legionella*-specific primer sets and quantitative techniques such as qPCR or ddPCR to directly measure the absolute abundance of *Legionella* in soils.

### Biogeography and climate structure global soil *Legionella* diversity

Multiple lines of evidence indicate that uncultured *Legionella* account for the vast majority of within-genus diversity (1,10,47), a conclusion reinforced by our observation that fewer than 2% of *Legionella* sequences matched ASVs corresponding to characterized species. This gap likely reflects limitations of cultivation approaches, as *Legionella* typically require specific growth conditions or host associations that are rarely reproduced *in vitro* (9,92–93). In addition, most isolation efforts and culture media have historically targeted clinical isolates or water-associated sources, leaving soil-associated species comparatively underexplored (15).

At a global scale, *Legionella* community similarity declined gradually across environmental gradients and geographic distance. This pattern is consistent with models of microbial biogeography in which environmental filtering and spatial processes jointly structure community composition (94). Furthermore, dominant lineages exhibited substantial regional differentiation, with the highest regional specificity occurring in geographically isolated areas such as Chile and Iceland. This pattern is consistent with studies of island and geographically isolated ecosystems, which suggest that dispersal limitation and historical contingency can influence which taxa become abundant (95–96). In addition, climatic conditions also shape which *Legionella* ASVs become regionally abundant: taxa were more frequently shared among regions with similar temperature and precipitation regimes, consistent with patterns resulting from environmental filtering (97). Dominant *Legionella* ASVs also showed phylogenetic clustering, indicated by high NRI and NTI values within several regions, further supporting homogeneous selection on phylogenetically conserved traits and resulting in assemblages dominated by closely related taxa (98).

A complementary explanation for the observed phylogenetic clustering is biotic filtering mediated by protist hosts. As *Legionella* replicate intracellularly within amoebae (9), host compatibility can be a major determinant of environmental persistence. If regional differences in amoebal community composition structure which *Legionella* lineages are able to proliferate, then local assemblages may reflect host-mediated selection acting within a broadly distributed evolutionary pool. Such dynamics could generate tip-level clustering without producing regionally endemic deep clades. Because global 18S rRNA gene datasets remain far less abundant than 16S datasets (90), and paired bacterial-protist datasets are still relatively scarce (99), the role of protist community structure in shaping *Legionella* biogeography remains underexplored and represents an important avenue for future research.

### Distribution of potentially pathogenic *Legionella* in soils

Pathogenic or potentially pathogenic ASVs were consistently detected globally, and their enrichment under warmer and wetter conditions aligns with epidemiological evidence linking legionellosis incidence to increased temperature and precipitation. As such, environmental reservoirs such as soils may represent an underappreciated exposure pathway, particularly under conditions where soil disturbance can generate aerosols. Notably, the most clinically significant species, *L. pneumophila* (1), was rare in soils, whereas the ASV corresponding to *L. longbeachae* (47), a species commonly associated with soil environments, ranked among the more abundant *Legionella* lineages detected. Indeed, the prominence of *L. longbeachae* in soils is consistent with epidemiological patterns in Australasia, where potting soils are a major source of infection (47). The enrichment of pathogenic lineages under warmer and wetter conditions is also consistent with the expectation that species with higher infectivity may be better adapted to elevated temperatures and may maintain closer evolutionary relationships with amoebal hosts, which serve as natural reservoirs and training grounds for intracellular pathogens (9,100). This underscores that infection risk is shaped not solely by environmental abundance but also by transmission pathways and genomic traits that are selectively favored in human-associated environments such as engineered water systems. Accordingly, understanding pathogenic mechanisms of environmentally-enriched *Legionella* taxa may be important under future climatic conditions if natural reservoirs become a significant source of transmission.

Comparisons of 16S rRNA genes from complete and reference-quality *Legionella* genomes demonstrates that the commonly sequenced 16S rRNA gene region provides moderate taxonomic resolution within the genus, resolving many species into unique ASVs while collapsing others - including several clinically relevant taxa - into shared sequence variants. While in this study we only utilized the first 150 bp of the 515F-806R region, additional analysis indicates that these trends would persist even when using the entire region (Fig. S14). This partial resolution underscores a key limitation of amplicon-based approaches for inferring pathogenic potential: the presence of an ASV matching a known pathogen does not necessarily uniquely identify a single pathogenic species. Additionally, the absence of such matches also does not preclude pathogenic capacity, given that pathogenicity in *Legionella* is narrowly defined by culture-confirmed human infections (101). Further studies are needed to evaluate *Legionella*-specific primer sets for taxonomic resolution, coverage and specificity. Additionally, since our analysis is based on amplicon sequencing of environmental DNA, detection does not necessarily indicate metabolically active populations. Future work incorporating RNA-based approaches, environmental aerosol measurements, or targeted cultivation may help distinguish active or potentially transmissible populations from dead or dormant cells.

### Concluding remarks

In summary, our global analysis parses large-scale microbiome datasets to show that *Legionella* are a widespread but low-abundance member of soil microbiomes whose distribution is strongly structured by climate and biogeography. The enrichment of *Legionella* and pathogenic *Legionella* ASVs under warmer and wetter conditions suggests that climatic factors play a critical role in shaping environmental reservoirs of these bacteria. Future studies using *Legionella*-specific primer sets and quantitative approaches such as qPCR or ddPCR will help resolve absolute abundance patterns and clarify the role of soils as potential reservoirs of exposure.

## Methods

### Global soil 16S rRNA gene amplicon datasets

Publicly available 16S rRNA gene amplicon datasets were retrieved from the NCBI Sequence Read Archive (SRA) (102). Studies were identified through targeted searches for soil microbiome datasets and either (i) spanned broad continental-scale geographic ranges or (ii) consisted of individual field sites that were manipulated to test specific environmental variables. To enable global-scale comparisons, only studies including at least the first 150 bp downstream of the bacterial 16S rRNA gene 515F primer were included (58–73). Studies were further screened to ensure the availability of geographic coordinates, sampling date, and key edaphic metadata, including soil pH, total organic carbon, and total organic nitrogen. Climatic variables, including midrange monthly temperature at the time of sampling and annual precipitation, were extracted from the WorldClim database using sample coordinates (78).

### Sequence processing

Processed 16S rRNA gene amplicon sequences in FASTA format were downloaded from the NCBI Sequence Read Archive (SRA) and subjected to a standardized downstream processing pipeline. Sequences were trimmed to a uniform length of 150 bp using seqkit (103), and reads shorter than 150 bp were removed. Quality-controlled sequences were denoised using Deblur (104), resulting in error-corrected amplicon sequence variants. Resulting sequences were dereplicated using VSEARCH (105) to obtain sequence abundances across samples. To reduce potential artifacts and sequencing noise, sequences were clustered at 99% identity and global singleton clusters were removed to reduce potential artifacts. After arriving at this final dataset, all downstream analyses were performed using amplicon sequence variants (ASVs) rather than clustered OTUs to get fine-scale resolution.

### Taxonomic identification of *Legionella*

To maximize sensitivity and specificity of *Legionella* detection, we implemented a two-step taxonomic classification approach. First, all ASVs were screened using BLAST (106) against a custom *Legionella* reference database. This database comprised 16S sequences from all reference-quality and complete *Legionella* genomes available in the NCBI database, together with all sequences from neighboring family Coxiellaceae (also within the order *Legionellales*), as well as a representative selection of 16S sequences spanning the broader Gammaproteobacteria and bacterial domain. Candidate sequences were then placed into a backbone phylogenetic tree constructed from this custom reference database using PhyML (107) to confirm phylogenetic affiliation. Only ASVs that clustered within the *Legionella* clade were retained. The accuracy and specificity of this approach was verified using all Gammaproteobacteria reference sequences from the SILVA database, as well as a mock community composed of DNA from *L. pneumophila*, *L. longbeachae* and *L. micdadei*.

### Association of *Legionella* relative abundance with environmental metadata

*Legionella* relative abundance was calculated as the proportion of *Legionella* reads relative to the total number of bacterial reads per sample. Samples lacking detectable *Legionella* sequences were retained for presence-absence and modeling analyses. Associations between *Legionella* relative abundance and environmental variables were assessed using Spearman rank correlations. For field experiments involving controlled environmental manipulations, *Legionella* relative abundance was compared between treatment and control plots using non-parametric tests or correlation analyses, as appropriate for each experimental design. To examine non-monotonic and threshold responses, each environmental variable was partitioned into five bins spanning the full observed range, and differences in *Legionella* relative abundance among bins were tested using Kruskal-Wallis tests followed by Dunn’s post hoc comparisons with multiple-testing correction. To evaluate interactive climatic effects, *Legionella* relative abundance was modeled across the joint temperature-precipitation space using a two-dimensional generalized additive model (GAM) (108). Model performance was compared to additive models with independent smooth terms for temperature and precipitation using analysis of variance (ANOVA) to assess improvement in fit.

### Mixed-effects modeling of *Legionella* relative abundance

To evaluate the joint effects of climatic and biogeochemical variables while accounting for study-level variation, a hurdle modeling framework was applied (109). Presence-absence of *Legionella* was modeled using a binomial mixed-effects model, and relative abundance conditional on presence was modeled using a beta mixed-effects regression. Study identity was included as a random intercept in both models to account for non-independence among samples (110). Predictor variables included temperature, precipitation, soil pH, elevation, total organic carbon, and total organic nitrogen.

### *Legionella* community composition and diversity

*Legionella* community composition was assessed using Bray-Curtis dissimilarities (111) calculated from *Legionella*-only ASV profiles. Ordination was performed using non-metric multidimensional scaling (NMDS) (112). Differences in community composition among regions were tested using permutation-based multivariate analysis of variance (PERMANOVA) (113). Environmental variables were fitted onto ordinations using permutation-based vector fitting (envfit) (114). Spatial structuring of *Legionella* communities was assessed by examining the relationship between Bray-Curtis similarity and geographic distance, with geographic distance calculated from sample coordinates. Distance-decay relationships were modeled using power-law regression (115). Multiple regression on distance matrices (MRM) was used to test whether geographic distance remained a significant predictor of community dissimilarity after accounting for climatic and biogeochemical differences among sites (116).

To examine dominant *Legionella* ASVs and their regional structuring, the 300 most abundant *Legionella* ASVs within each geographic region were selected. Overlap among regions were quantified using pairwise comparisons, and relationship between ASV overlap and climatic dissimilarity among regions were assessed using Spearman correlations. To evaluate whether biogeographic and climatic regions displayed deep phylogenetic divergence, a maximum-likelihood phylogeny was constructed using the 300 most abundant *Legionella* ASVs from each region together with reference sequences from complete and reference-quality *Legionella* genomes. Sequences were aligned using MAFFT (117), and phylogenetic inference was performed using IQ-Tree (118) under an appropriate nucleotide substitution model. Phylogenetic trees were visualized and annotated using iTOL (119). To quantify within-region phylogenetic structure, clustering and dispersion were assessed using the Net Relatedness Index (NRI) and Nearest Taxon Index (NTI) (98), calculated using the picante R package (120) as standardized effect sizes of mean pairwise phylogenetic distance (MPD) and mean nearest taxon distance (MNTD) relative to a null model with 999 randomizations. Significance was determined using rank-based two-tailed p-values (ɑ=0.01), where p <0.01 indicated significant clustering and p>0.99 indicated significant overdispersion.

### Community context of *Legionella*-rich samples

To identify microbial taxa associated with *Legionella*-rich communities, differential abundance analysis was performed using 97% OTUs, which were clustered using VSEARCH (105). Samples were stratified into upper and lower quartiles of *Legionella* relative abundance. Differentially abundant OTUs were identified using log_2_fold-change thresholds and false discovery rate (FDR) correction (121). Taxonomic composition and prevalence of enriched OTUs were plotted to assess taxa that were abundant in both *Legionella*-rich and *Legionella*-poor communities.

### Identification of pathogenic *Legionella* ASVs

Reference-quality and complete *Legionella* renomes were retrieved from NCBI (122), trimmed to the first 150 bp following the 515F primer and clustered to determine species-level resolution. ASVs matching reference sequences from known pathogenic species were designated as pathogenic ASVs, while ASVs shared with both pathogenic and non-pathogenic species were designated as potentially pathogenic. Associations between pathogenic or potentially pathogenic *Legionella* ASVs and environmental variables were assessed using logistic regression models. Presence-absence of pathogenic ASVs was modeled as a function of climatic and biogeochemical variables, with sequencing depth included as a covariate to account for detection bias.

## Acknowledgements

Research reported in this publication was supported by the National Institute Of General Medical Sciences of the National Institutes of Health under Award Number P20GM125508. The content is solely the responsibility of the authors and does not necessarily represent the official views of the National Institutes of Health. We thank the investigators and research teams who generated and made publicly available the sequencing datasets used in this study. We also thank the Center for Microbial Analysis, Innovation, and Knowledge Integration (CMAIKI) at the University of Hawai’i for computational resources and support.

## Data availability

All sequencing data used in this study are publicly available through the NCBI BioProject repositories listed in Supplementary Table 1. All analyses were conducted using open-access software.

---

**Fig. S1.**
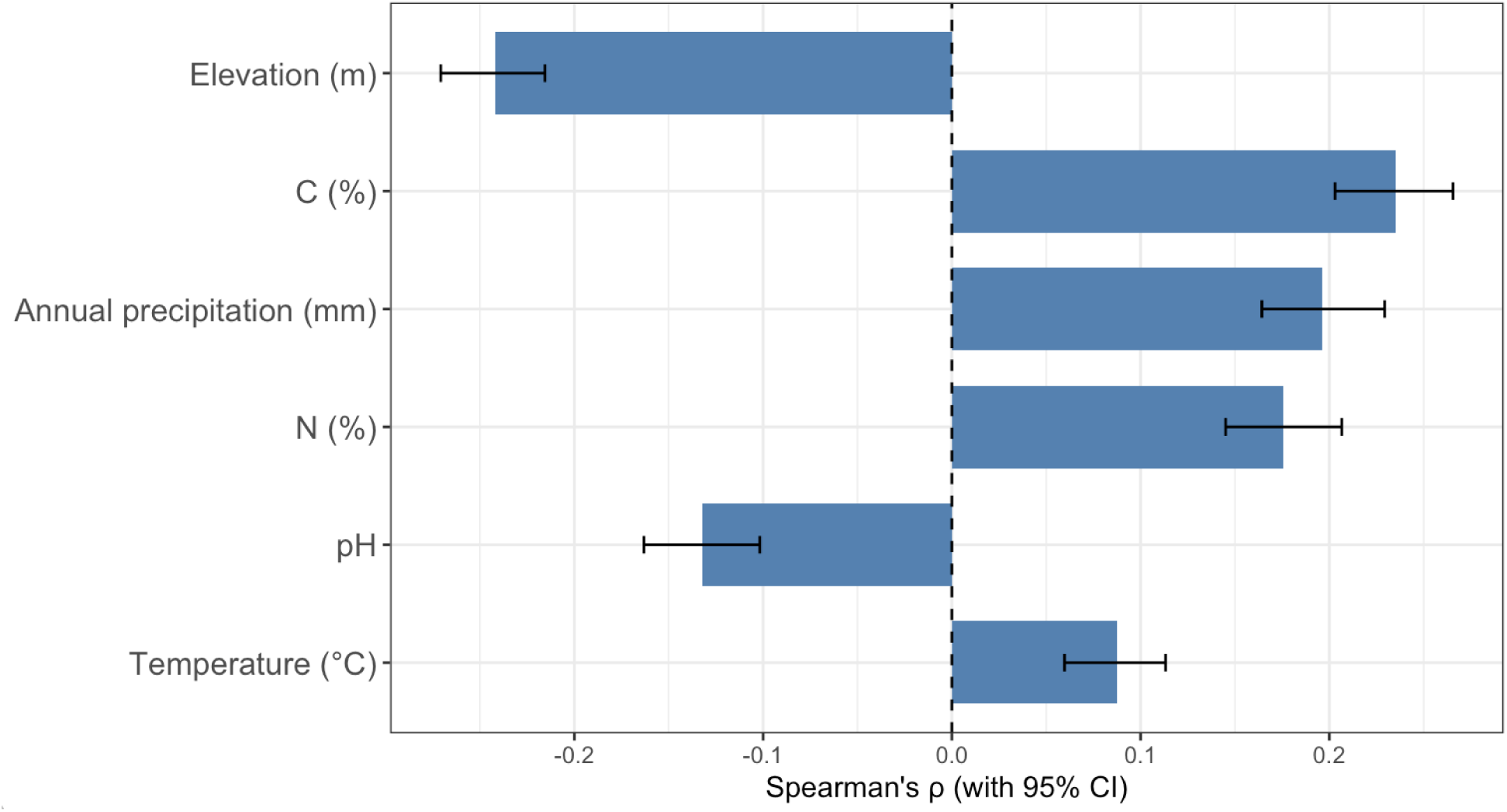
Spearman correlations between *Legionella* relative abundance and climatic and biogeochemical variables. Bars represent Spearman’s rank correlation coefficients (ρ) between *Legionella* relative abundance and each environmental variable across all samples. Error bars indicate 95% confidence intervals.

**Fig. S2.**
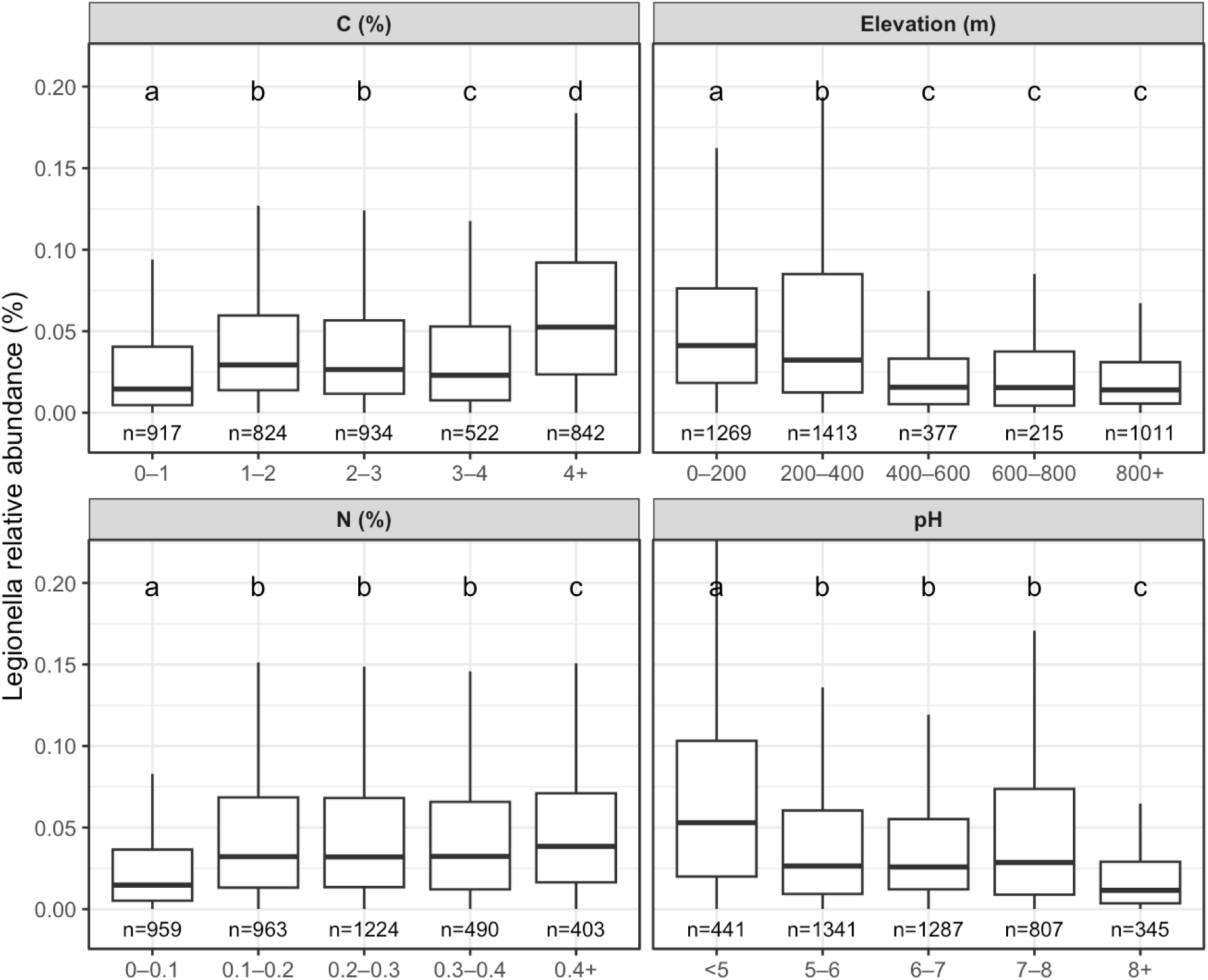
Variation in *Legionella* relative abundance across biogeochemical and abiotic gradients. Boxplots show *Legionella* relative abundance across five bins spanning the observed range of soil carbon (C), elevation, nitrogen (N) and pH values. Boxes represent the interquartile range (IQR), center lines indicate medians and whiskers extend to 1.5x the IQR. Sample sizes for each bin are indicated below the x-axis. Letters above boxes denote statistically distinct groups identified using Dunn’s post hoc comparisons following Kruskal-Wallis tests (p<0.05).

**Fig. S3.**
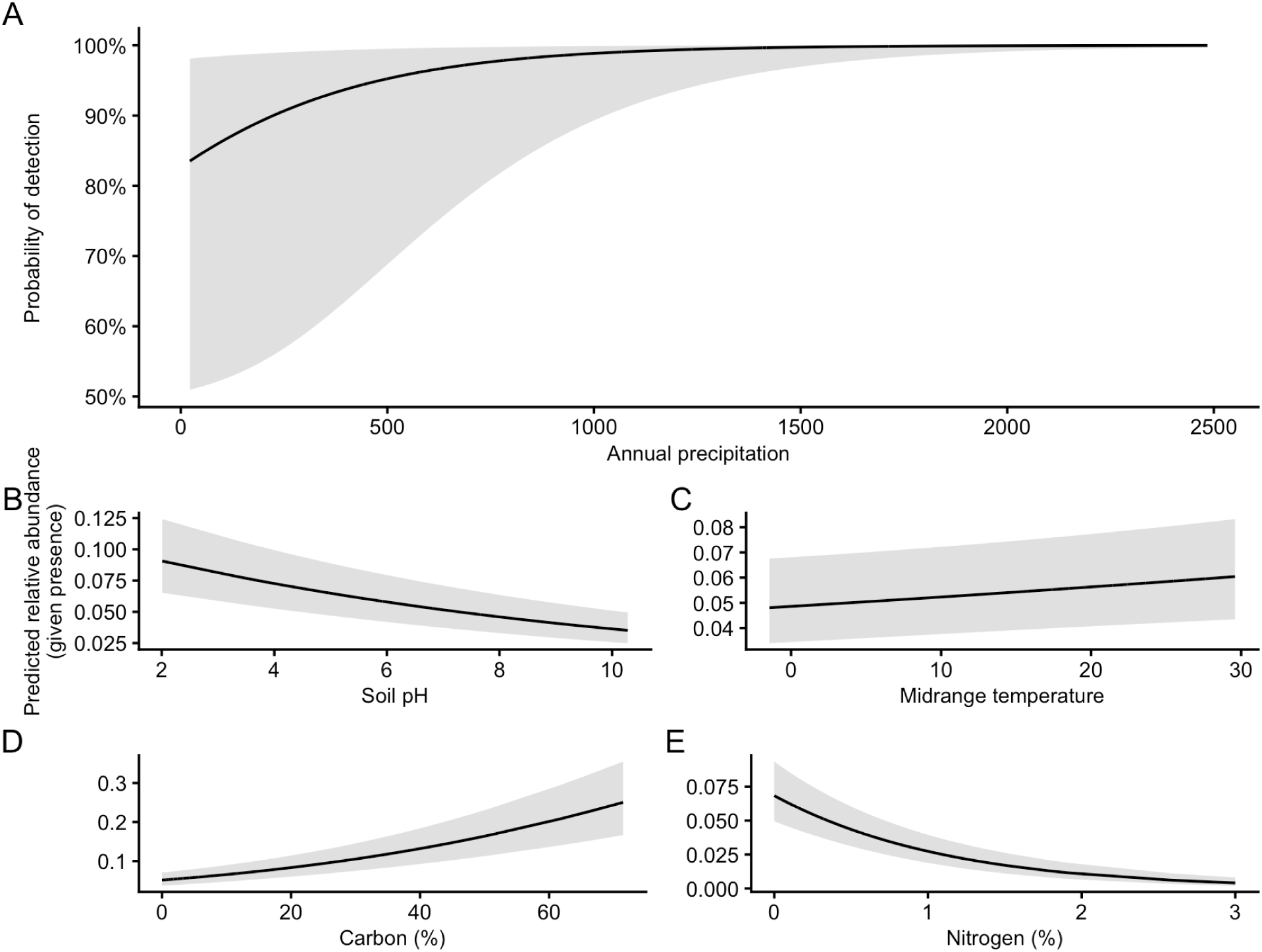
Hurdle mixed-effects model results for predictors of *Legionella* occurrence and abundance. (A) Predicted probability of detecting *Legionella* as a function of annual precipitation (mm) from a binomial mixed-effects model. (B-E) Predicted relative abundance of *Legionella* conditional on presence from a beta mixed-effects regression model across gradients of soil pH, midrange temperature, soil carbon (C), and soil nitrogen (N). Lines represent model predictions and shaded regions indicate 95% confidence intervals. Study was included as a random intercept in all models.

**Fig. S4.**
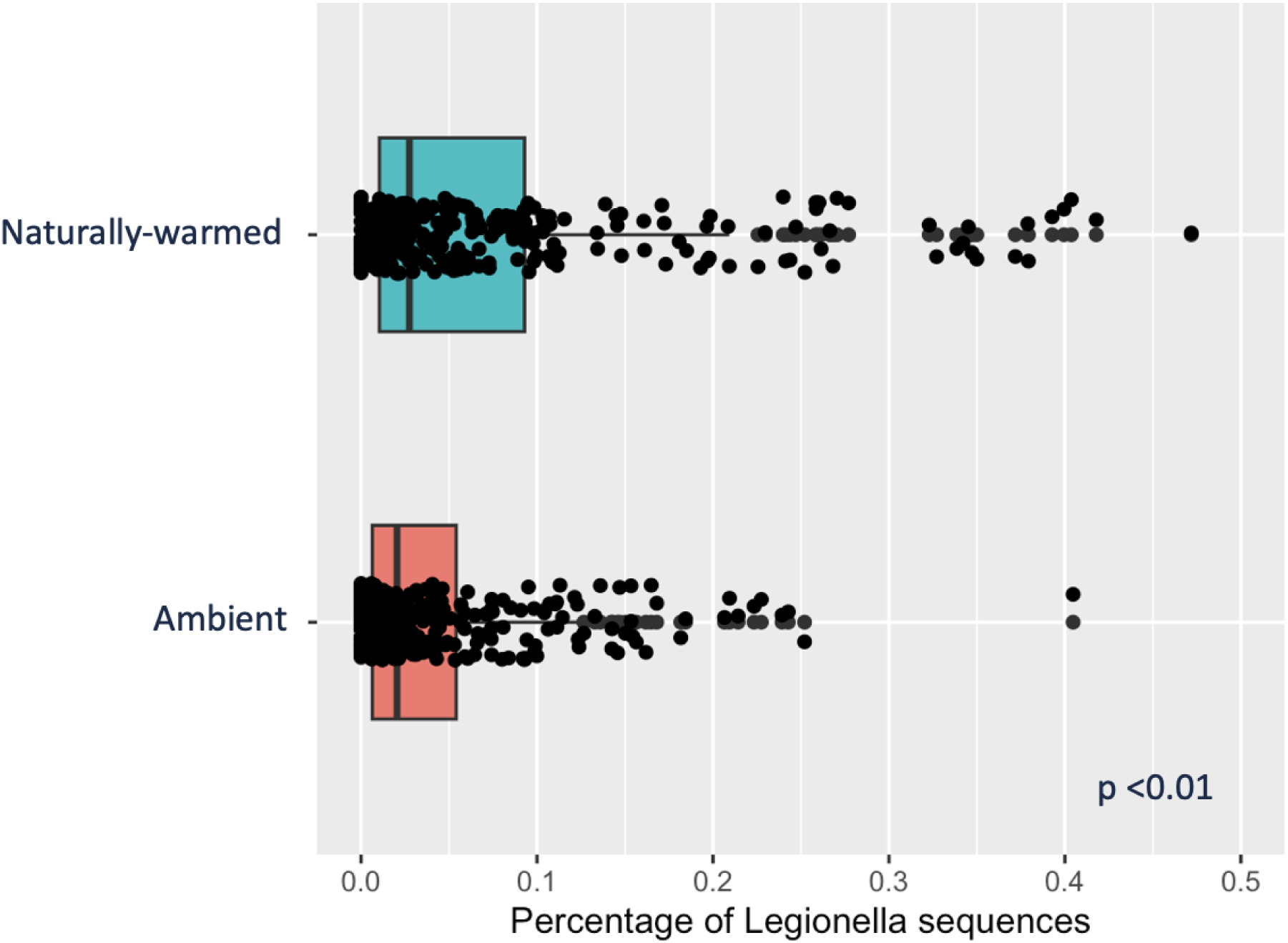
*Legionella* relative abundance at geothermally warmed and ambient sites in Iceland. Comparison of *Legionella* relative abundance between naturally warmed soil (+6°C) and nearby ambient sites. Points represent individual samples and boxplots summarize the distribution (median and interquartile range).

**Fig. S5.**
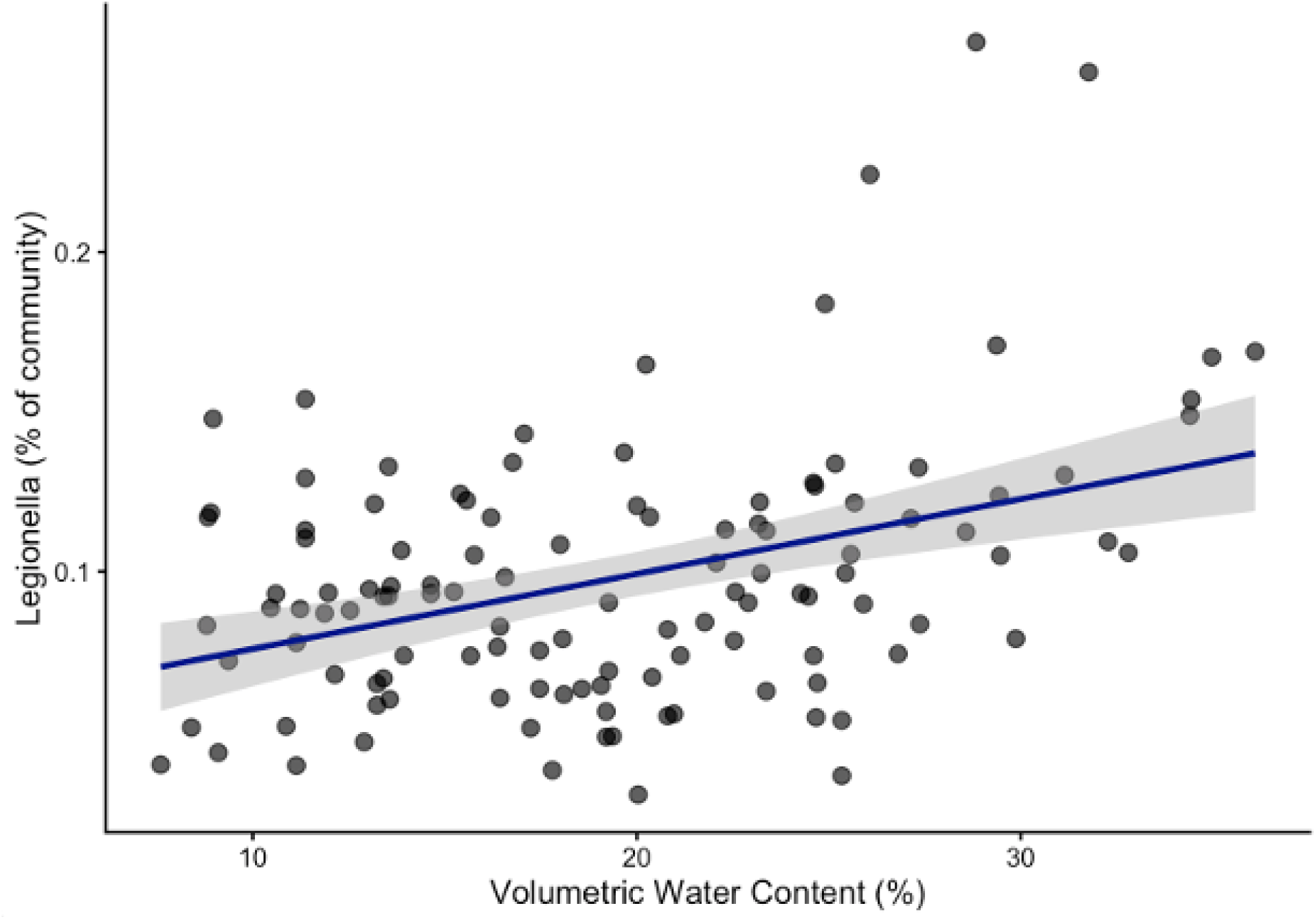
Relationship between soil moisture and *Legionella* relative abundance at rainfall-manipulation sites in Wisconsin. Relative abundance of *Legionella* sequences plotted against volumetric water content (%). Points represent individual samples. The line indicates the fitted linear regression and the shaded region shows the 95% confidence interval.

**Fig. S6.**
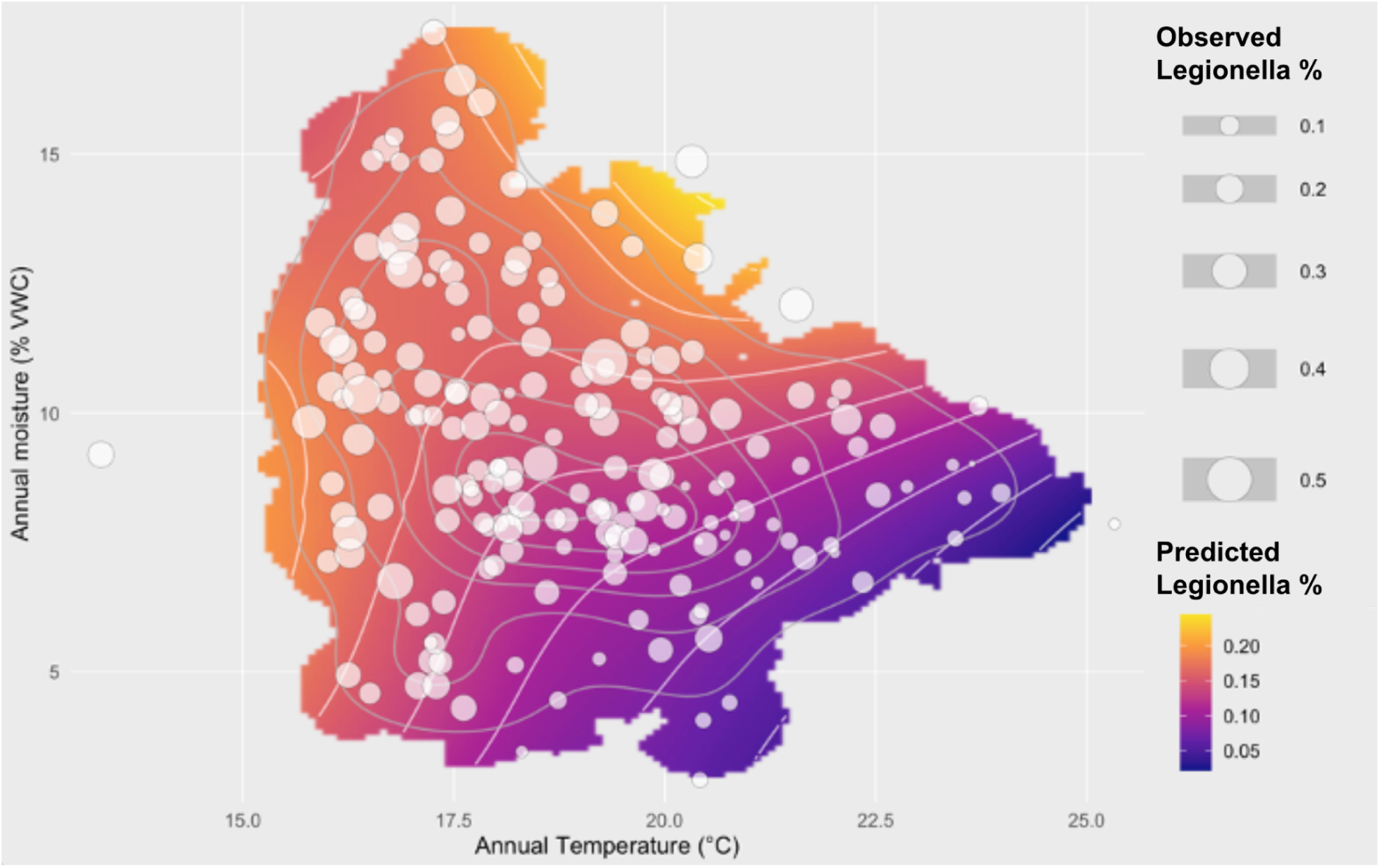
*Legionella* relative abundance across temperature and soil moisture gradients at a combined climate-manipulation experiment in Oklahoma. Observed *Legionella* relative abundance is shown as points, with point size proportional to the percentage of *Legionella* sequences detected in each sample. The colored surface represents LOESS-predicted *Legionella* relative abundance, with contour lines indicating predicted abundance levels.

**Fig. S7.**
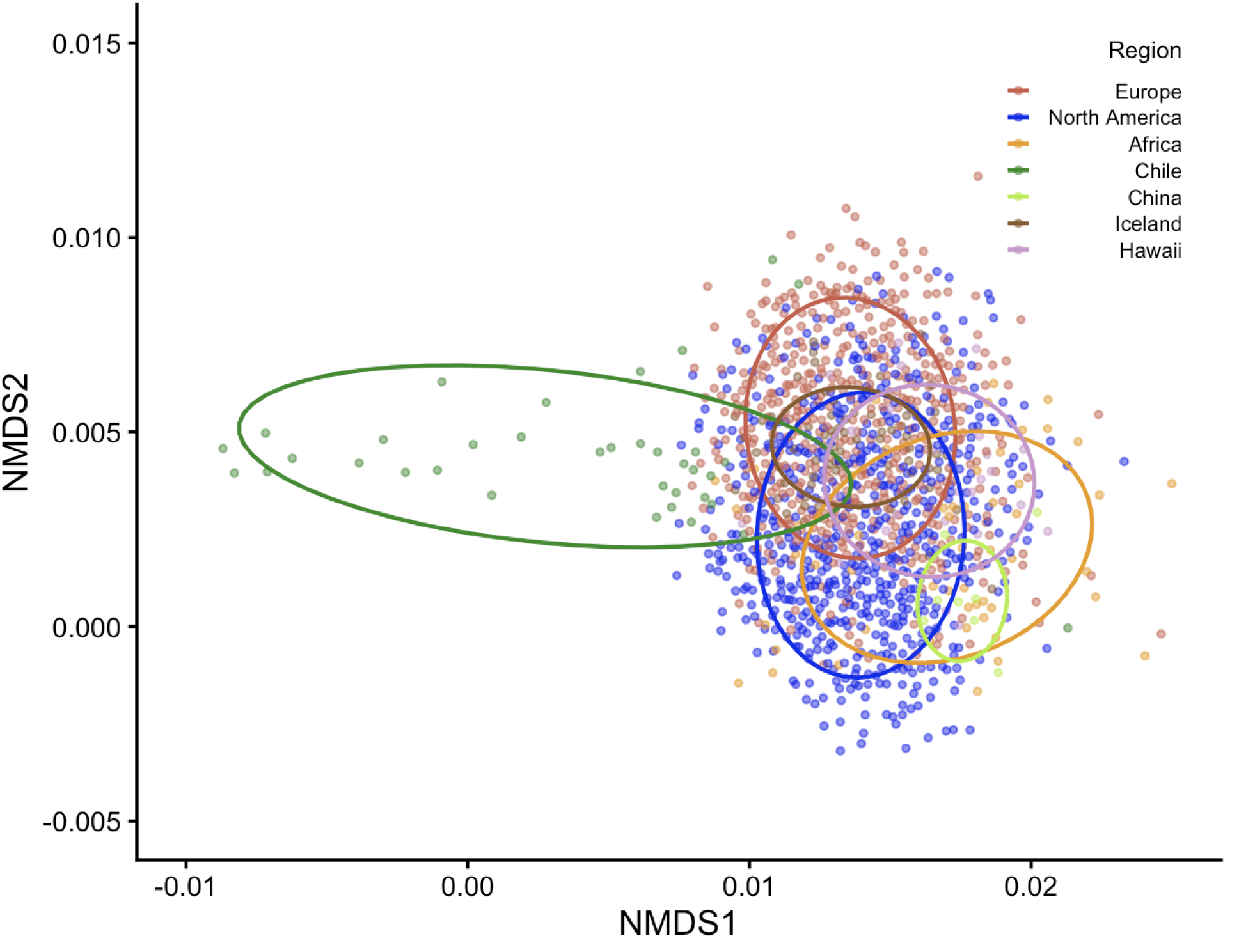
Regional structuring of *Legionella* community composition. Non-metric dimensional scaling (NMDS) ordination of *Legionella* community composition based on Bray-Curtis dissimilarities calculated from ASV relative abundances. Each point represents an individual sample and colors indicate geographic region. Ellipses represent 95% confidence intervals around the centroid of each region.

**Fig. S8.**
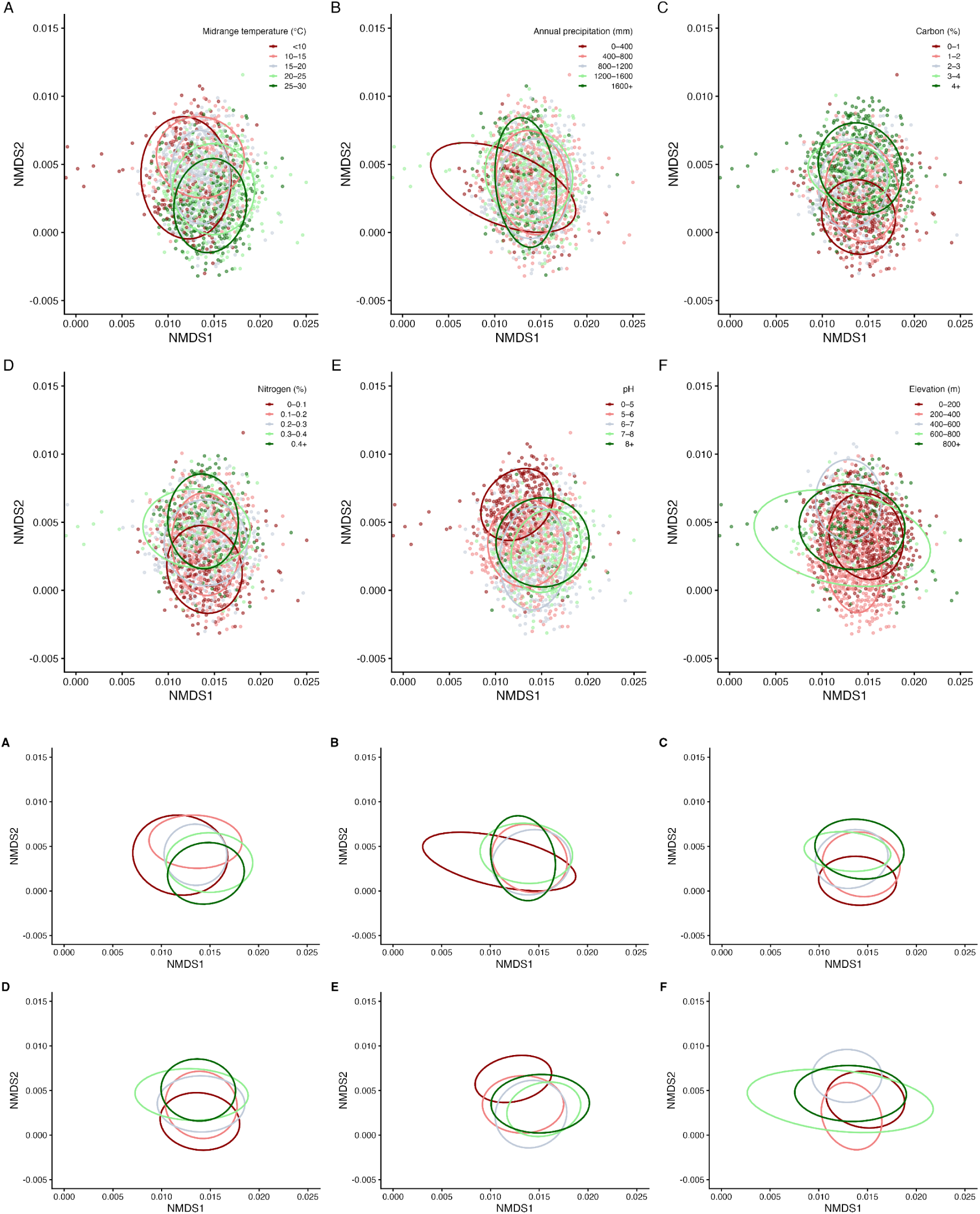
Environmental structuring of *Legionella* community composition. NMDS ordination of *Legionella* community composition based on Bray-Curtis dissimilarities calculated from *Legionella* ASV relative abundances. Samples are colored according to environmental bins for midrange temperature, annual precipitation, soil carbon (C), soil nitrogen (N), soil pH and elevation. Ellipses represent 95% confidence intervals around the centroid of samples within each environmental bin.

**Fig. S9.**
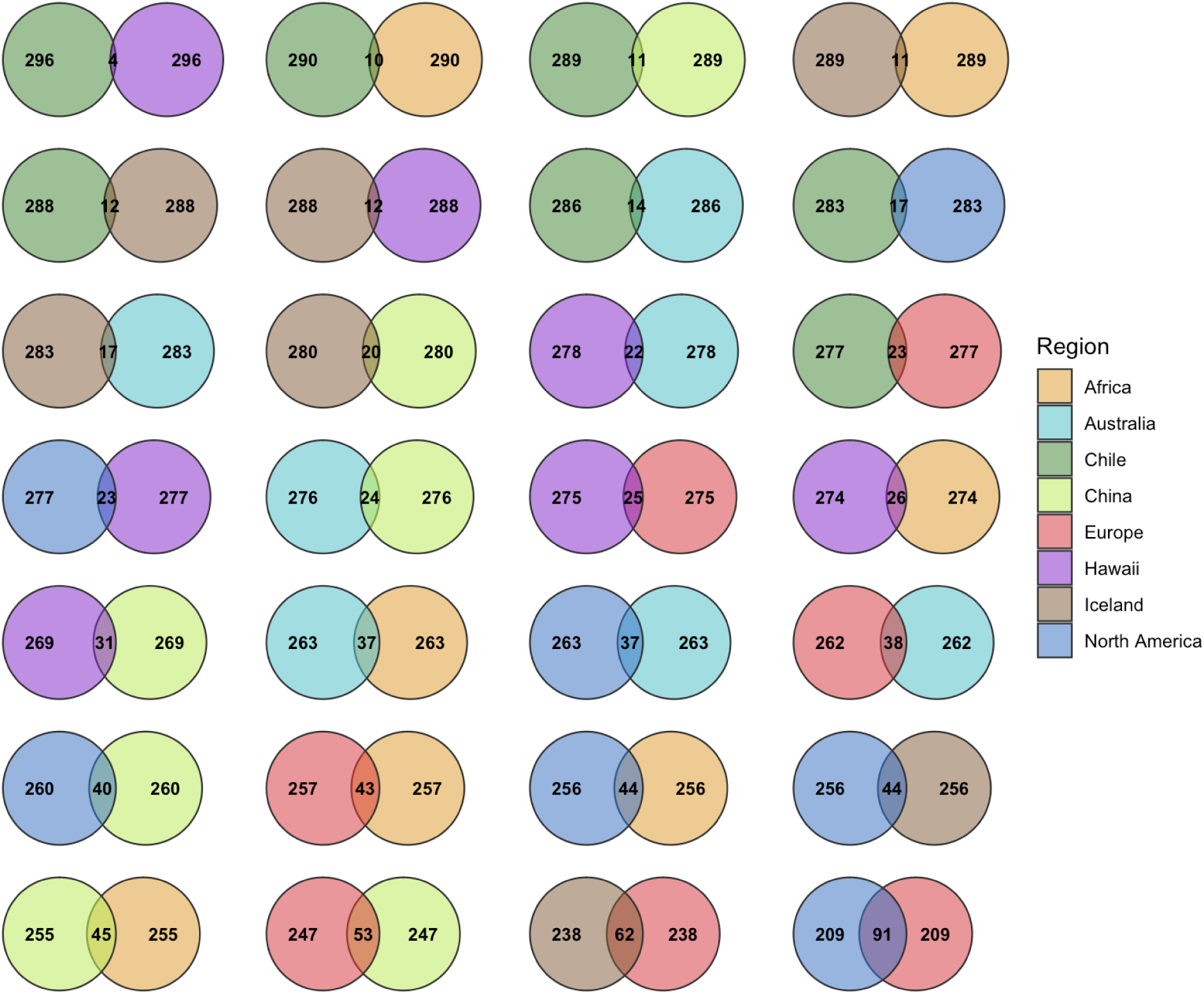
Pairwise overlap of dominant *Legionella* ASVs across regions. Venn diagrams illustrate overlap among the 300 most abundant *Legionella* ASVs for each pairwise comparison of the eight regions. Colors indicate regions, and numbers represent the number of dominant ASVs unique to each region (outer sections) or shared between regions (center). Comparisons are arranged from lowest overlap (upper left) to highest overlap (bottom right).

**Fig. S10.**
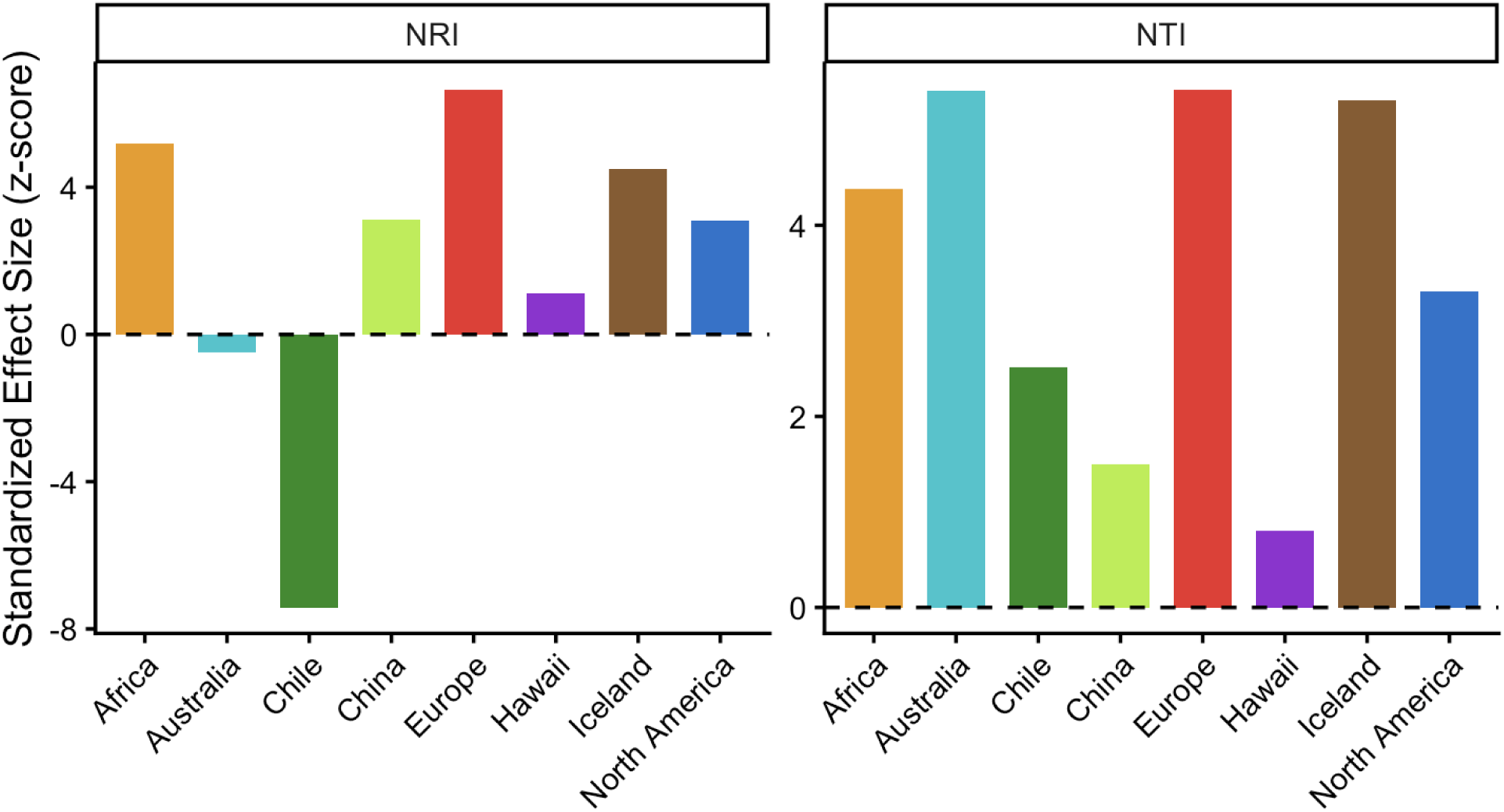
Phylogenetic structure of abundant *Legionella* ASVs across global regions. Net Relatedness Index (NRI) and Nearest Taxon Index (NTI) were calculated as standardized effect sizes (SES z-scores) relative to null model communities. Positive values indicate phylogenetic clustering, whereas negative values indicate phylogenetic overdispersion. Bars are colored by geographic region.

**Fig. S11.**
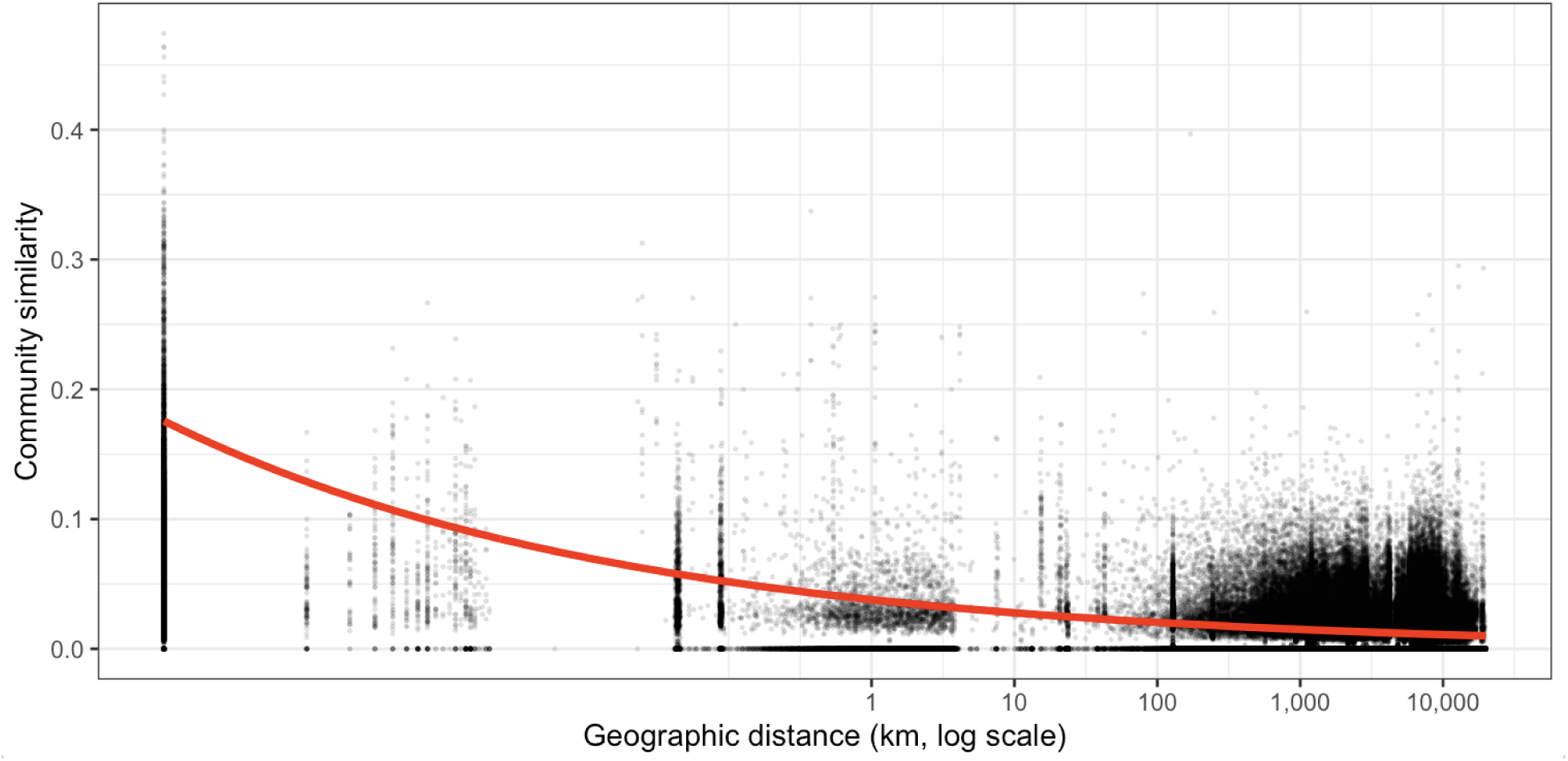
Distance-decay relationship of *Legionella* community composition. Pairwise community similarity among samples (1-Bray-Curtis dissimilarity) is plotted against geographic distance (km). Points represent pairwise comparisons between samples, and the red line indicates the fitted distance-decay relationship.

**Fig. S12.**
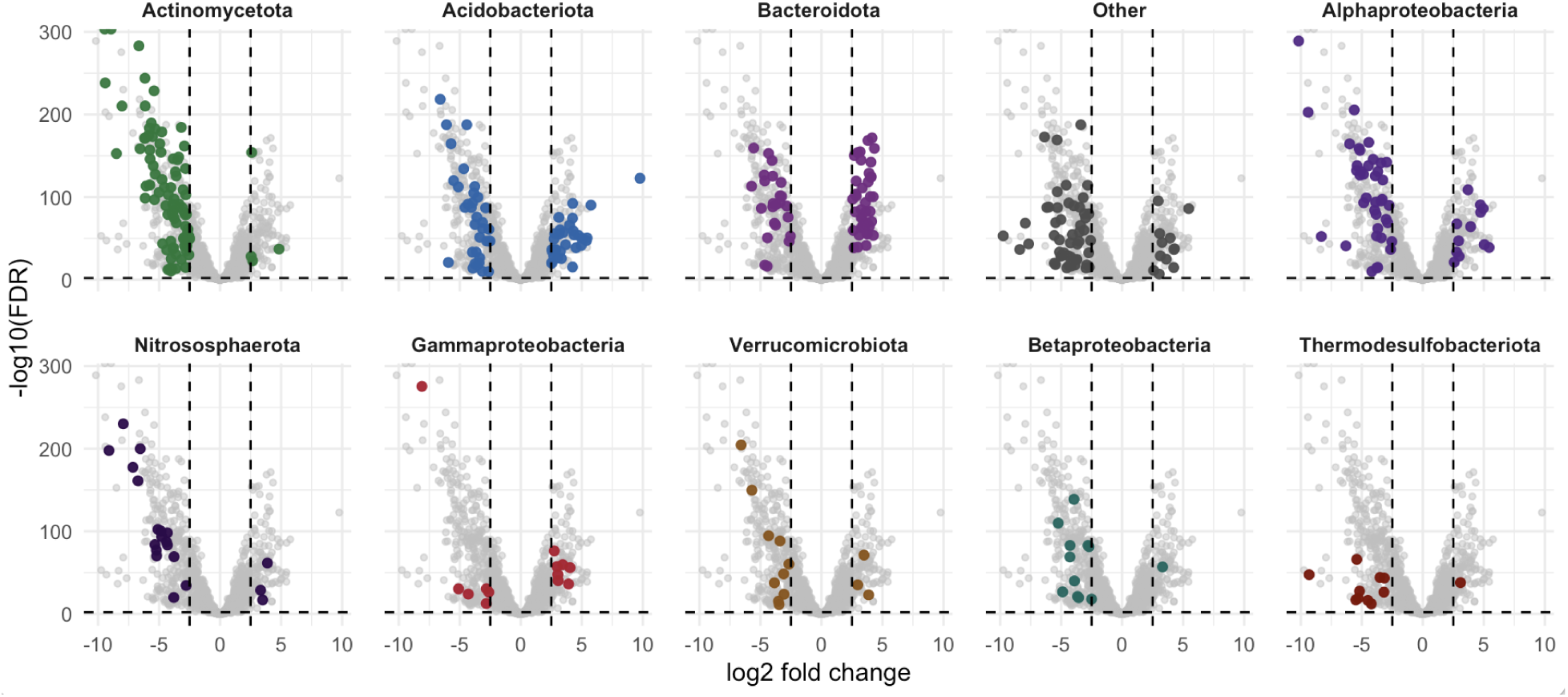
Differentially abundant microbial taxa associated with *Legionella* abundance. Volcano plot showing microbial community features (97% OTUs) enriched in *Legionella*-rich versus *Legionella*-poor samples. Samples were classified into upper (>0.062%) and lower (<0.009%) quartiles of *Legionella* relative abundance. Points represent OTUs with log₂ fold change on the x-axis and -log_10_(FDR) on the y-axis. Positive fold changes indicate enrichment in *Legionella*-rich samples, whereas negative fold changes indicate enrichment in *Legionella*-poor samples. Panels show major taxonomic groups, and colored points indicate OTUs significantly enriched in either direction (|log₂ fold change| ≥ 2.5, FDR < 0.01).

**Fig. S13.**
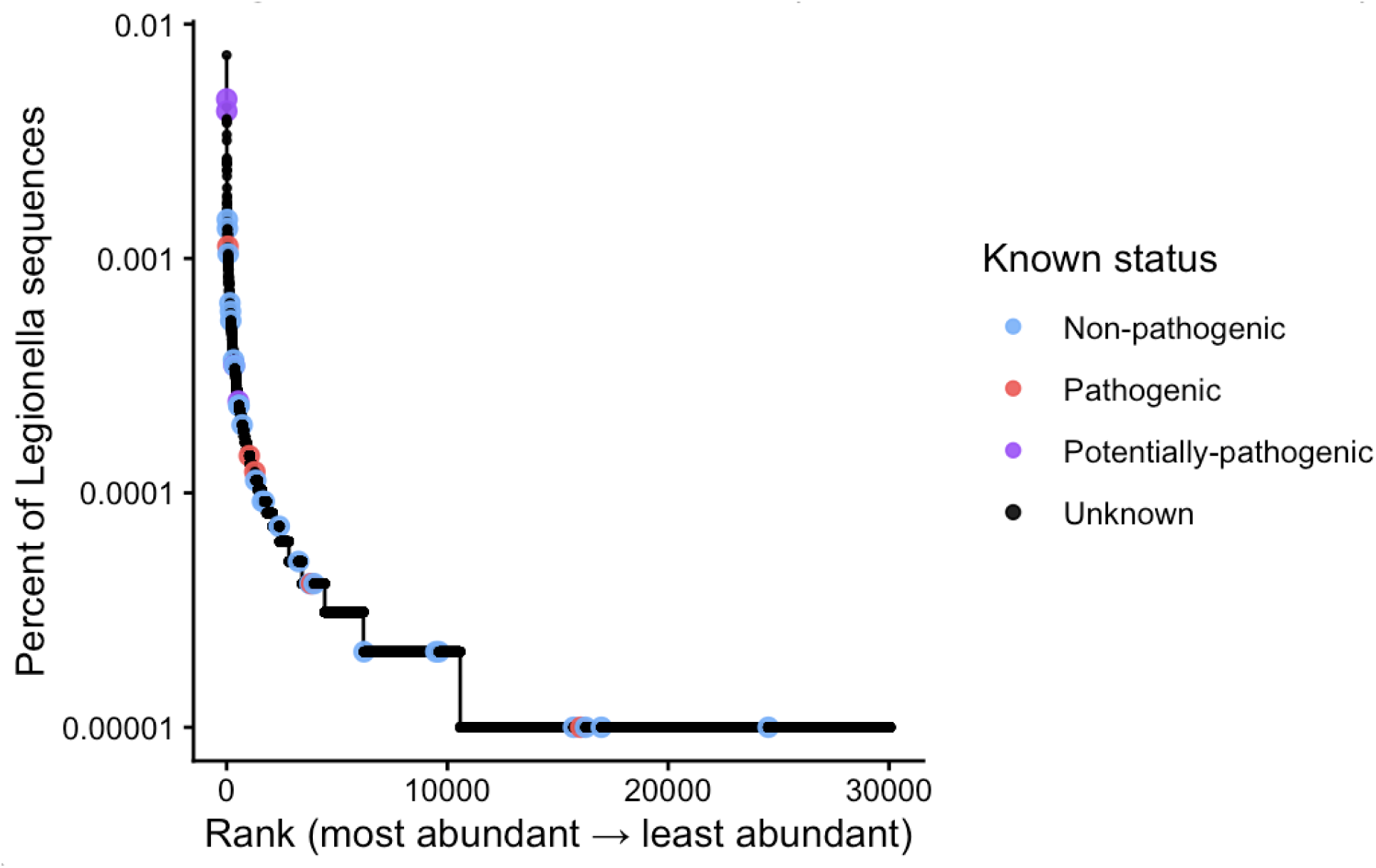
Rank abundance distribution of *Legionella* ASVs. Rank abundance plot showing the relative abundance of all *Legionella* ASVs recovered. ASVs are ordered from most abundant to least abundant along the x-axis, and the y-axis indicates the percentage of total *Legionella* sequences represented by each ASV. Points are colored according to pathogenic status based on matches to characterized species (non-pathogenic, pathogenic, potentially pathogenic, or unknown).

**Fig. S14.**
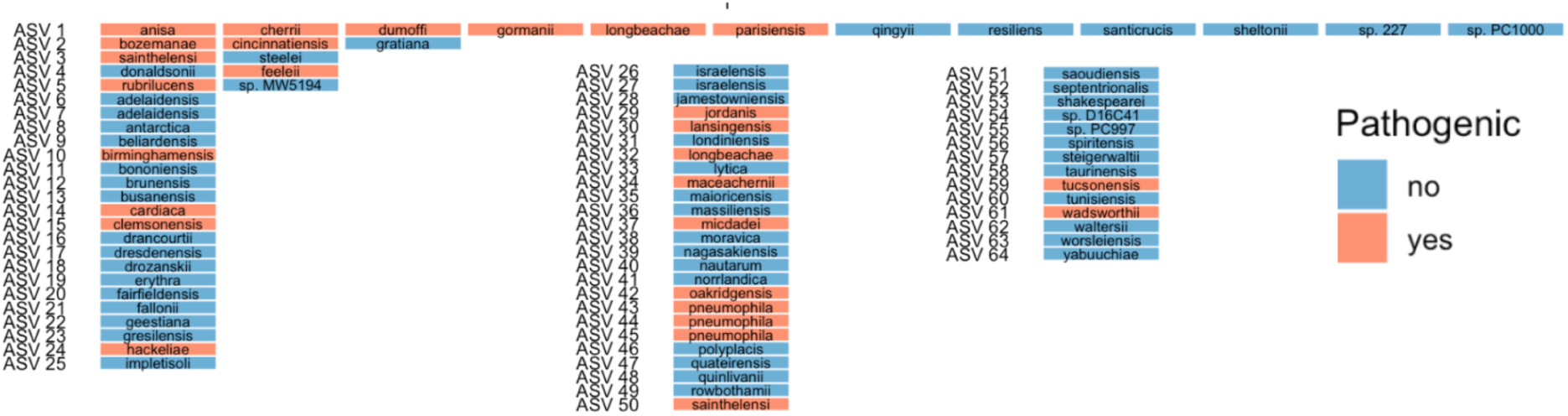
Clustering of characterized *Legionella* species based on 16S rRNA gene sequence identity. Clustering of characterized *Legionella* species based on the full 515F-806R 16S rRNA gene amplicon region. Species sharing identical sequences across this region are grouped into the same ASV. Boxes are colored according to whether the species is a known human pathogen.

